# Primate deep conserved noncoding sequences and non-coding RNA: their possible relatedness to brain and Central Nervous System

**DOI:** 10.1101/2021.08.17.456625

**Authors:** Nilmini Hettiarachchi

## Abstract

**Background:** Conserved non coding Sequences (CNSs) are extensively studied for their regulatory properties and functional importance to organisms. Many features such as location, proximity to the likely target gene, lineage specificity, functionality of likely target genes, and nucleotide composition of these sequences have been investigated, thus have provided very meaningful insight to signify underlying evolutionary importance of these elements. Also thorough investigation around how to assign function to non-coding regions of eukaryote genomes is another area that is studied. On one hand evolutionary analyses, including signatures of selection or conservation which can indicate the presence of constraint, suggesting that sequences that are evolving non-neutrally are candidates for functionality. On the other hand evidence that is based on experimental profiling of transcription, methylation, histone modifications and chromatin state. While these types of data are very important and are associated with function in most cases, this is not always the case. Evolutionary conservation though highly conservative which mostly considers elements identifiable in more than one species, is still being used as the initial guideline in investigating function via experiments. If we had an understanding of the experimental profiles of conserved non-coding regions as there may be patterns that are often associated these potentially functional elements it may help to construed functionality of conserved non coding regions easily.

**Results:** In an effort to try integrate experimental profile data, we investigated evidence of expression of conserved noncoding sequences (CNSs). For CNSs from ten primates, we assessed transcription, histone modifications, level of evolutionary constraint or accelerated evolution, and assessed possible target genes, tissue expression profiles of likely target genes (as some CNSs may be enhancers, and may be ncRNAs that interact directly with mRNA) and clustering patterns of CNSs. In total we found 153475 CNSs conserved across all ten primates. Of these 59,870 were overlapping non coding regions of ncRNA genes. H3K4Me1 marks (often associated with active enhancers) were highly correlated with CNSs whereas H4K20Me1 (linked to, e.g. DNA damage repair) had high correlation with conserved ncRNA regions (ncRNA-gene-CEs). Both CNSs and conserved ncRNA showed evidence of being under purifying selection. The CNSs in our dataset overall exhibited lower allele frequencies, consistent with higher levels of evolutionary constraint. We also found that CNSs and ncRNA-gene-CEs produce mutually exclusive groups. The analyses also suggest that both types of conserved elements have undergone waves of accelerated evolution, which we speculate may indicate changes in regulatory requirements following divergence events. Finally, we find that likely target genes for hominoidae, primate and mammalian-specific CNSs and ncRNA-gene-CEs are predominantly associated with brain-related function in humans.

**Conclusion:** The deep conserved primate CNSs and ncRNA gene-CEs signify functional importance suggesting ongoing recruitment of these elements into brain-related functions, consistent with King and Wilson’s hypothesis that regulatory changes may account for rapid changes in phenotype among primates.

## Introduction

Conservation in the non-coding regions of genomes are elaborately studied over more than a decade and still it keeps revealing intriguing information about the non-coding regions of genomes, which we once considered as “junk” DNA. It is very clear now that once referred to as “junk” shapes the lineage and species individuality.

Assessing the functional potential of non-protein coding regions of genomes is a tasking process. Two broad approaches focus on either evolutionary signals of conservation and non-neutrality (Sandelin et al. 2004;Woolfe et al. 2005; Vavouri et al. 2007; Elgar 2009; Lee et al. 2011;Matsunami and Saitou 2012; Takahashi and Saitou 2012;Hettiarachchi et al. 2014; Hettiarachchi et al. 2016) or ‘activity’ signals, such as expression, epigenetic or chromatin states. These approaches have led to strikingly different conclusions regarding the extent of the genome which is functional. Evolutionary analyses estimate that ∼10% of the genome is under constraint, whereas biochemical activities have been attributed to ∼80% of the human genome. This has led to major debate regarding the identification of biological function. On the one hand, it is clear that evolutionary analyses may be conservative, in that they frequently rely on conservation of an element in more than one species in order to predict function. Thus, they may miss species-specific elements or activities. In contrast, a difficulty with assessing functional regions through measurement of biochemical activities is that it is not clear how to separate bona fide functional activities from background noise. While there are good theoretical reasons to expect noise in the genomes of multicellular eukaryotes with small effective population size, a baseline has yet to be established, and may be non-trivial to achieve in practice.

Clearly both datasets are useful, but the debate that followed the publication of the ENCODE project (The ENCODE Project Consortium 2012) suggests that a cautious approach to extending evolutionary analyses to include biochemical activity data is warranted (Graur et al. 2013; Doolittle 2013). In an attempt to understand whether biochemical activity data relate to evolutionarily-derived functional annotation, we here integrate some of these data into an analysis of conserved non-coding elements from primates. We used expression data to divide these into CNSs (where ncRNAs are both conserved and expressed), and find that distinct types of histone modification are associated with CNSs and ncRNAs. For CNSs, it is thought that these may act on adjacent protein-coding genes (Woolfe et al. 2005; Elgar 2009; Matsunami and Saitou 2012), whereas the trans-acting nature of ncRNAs suggests this need not be the case for this class of non-coding element. We nevertheless applied this criterion, as some enhancers show evidence of transcription (Andersson et al. 2014; Melamed et al. 2016; Tippens et al. 2018), blurring the lines between ncRNAs and cis-acting regulatory elements.

Lastly, we found that our primate non-coding elements were most frequently associated with protein-coding genes involved in brain development. This suggests that location is non-random, and may further help in assessing non-coding element function. While we would be hesitant to assign function based on the non-evolutionary components of our analyses (histone modification, genomic location), these may nevertheless be helpful parameters in function of genomic elements where comparative data are absent.

## Materials and Methods

### Genomes used in the analysis

Primate repeat masked genomes of human (*Homo sapiensapien*), Gorilla ( *Gorilla gorilla gorilla*),Orangutan (*Pongoabelii*),Gibbon (*Nomascusleucogenys*), Rhesus macaque (*Macacamulatta*),M.fascicularis (*Macacafascicularis*),Baboon (*Papioanubis*),Vervet (*Chlorocebussabaeus*),Marmoset (*Callithrixjacchus*),Mouse lemur (*Microcebusmurinus*)) were downloaded from Ensembl release 94

All genomes used were above minimum of 5X coverage.

Figure 1 depicts the phylogenetic relationships between the species used in the study. The divergence times used for the work are according to Goodman et al. 1998;Glazko and Nei. 2003.

**Figure 1:**
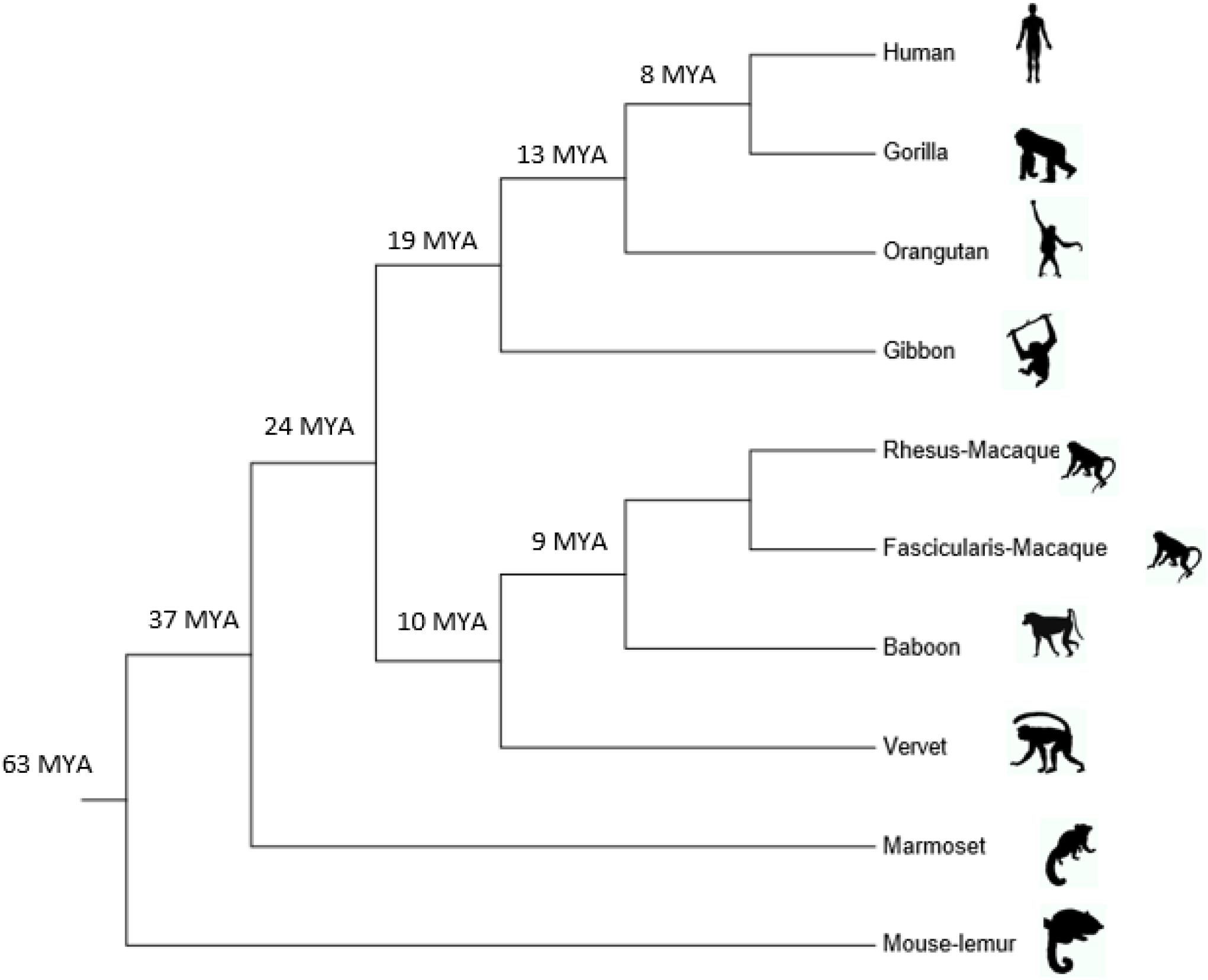
Phylogenetic relationships between the primate species used in the study. (MYA – Million Years Ago). The data for these species were downloaded from Ensembl release 94. The divergence times are according to Goodman et al. 1998; Glazko and Nei. 2003.

**Figure 2:**
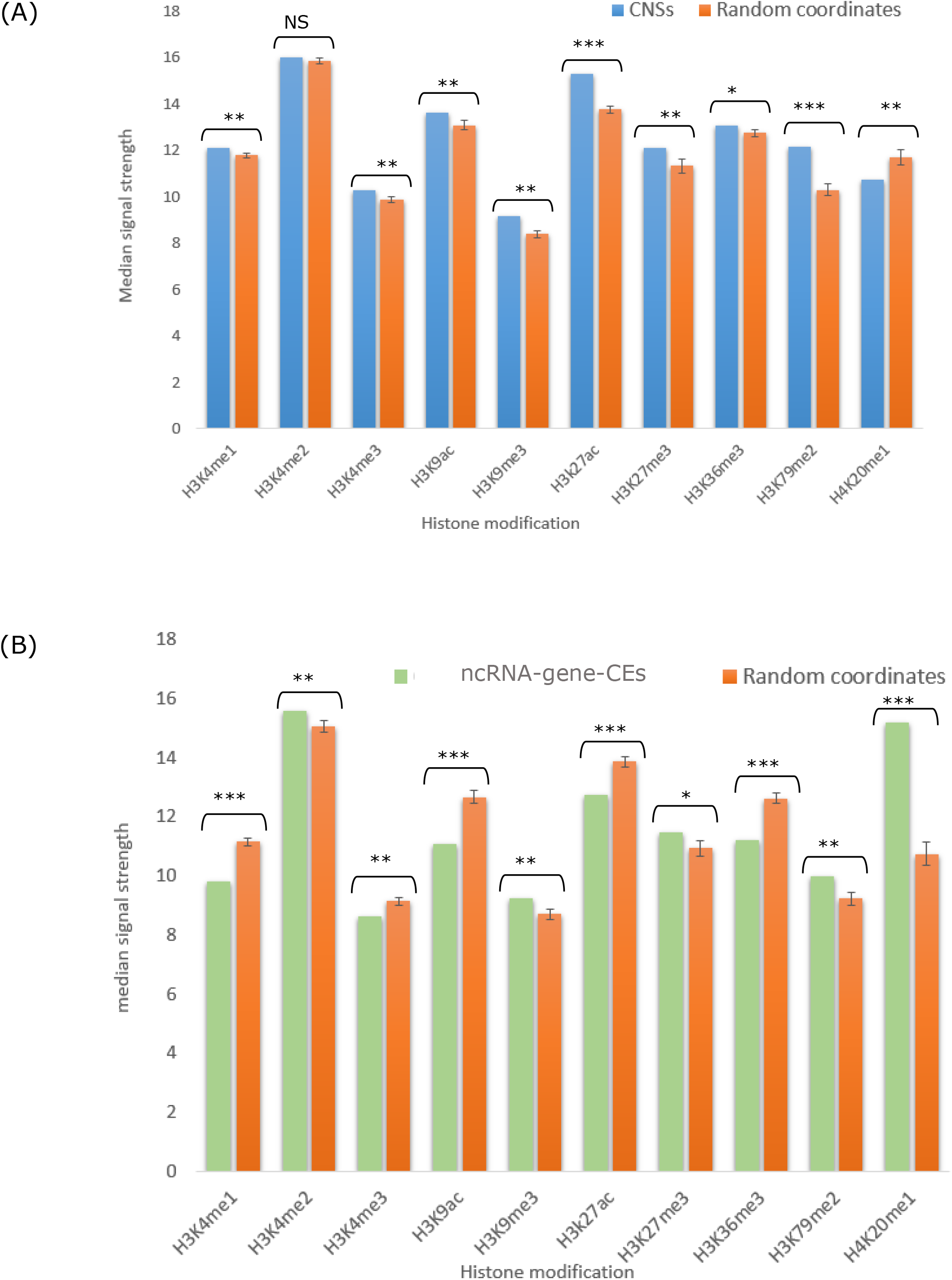
(A) Histone modification median signal strength for primate common CNSs. (B) Histone modification median signal strength for primate common ncRNA-gene-CEs. Specifically CNSs show higher signal strength for H3K27ac (histone marks associated with active enhancer regions) compared to conserved ncRNA and also random expectation. Conserved non-coding RNA shows significantly higher signal strength for H4K20me1. Statistical significance was determined via one-sample t-test. (P <0.0001 - ***, P > 0.001 = **, P>0.01 = *)

### Masking coding regions

CDS coordinates were downloaded for respective genomes from EnsemblBioMart release 94. The CDS coordinates of different transcripts were merged to remove overlapping or redundant coordinates. If a pair of coordinates were overlapping they were merged into one continuous length. The merged CDs coordinates were masked in the respective genomes, which results in only non-coding genomes.

### Homology search

After masking coding regions we searched for genomic locations that are conserved across all 10 primate species used in the study. BlastN 2.6.0 (Altschul et al. 1997) was used for determining homologous regions in a whole genome chain search starting from the most closely related species pair used in the study (Human and Gorilla). Mitochondrial DNA was removed from the analysis.

The commonly found conserved regions that exceeded 1:1 (one to one) protein coding threshold determined between human and Mouse lemur in all the 10 species were considered as primate common CNSs.

The determined commonly conserved sequences were classified into two groups based on how many conserved regions overlapped with annotated non-coding RNA gene regions in human. The annotation data for ncRNA genes is based on Ensembl version 94. Any conserved noncoding element that overlapped with annotated ncRNA genes were classified as primate common ncRNA-gene-CEs and remaining sequences are referred to as CNSs (Conserved Noncoding Sequences) hereon.

### Determining histone modification and DNaseI hypersensitive site overlaps for primate common CNSs and ncRNA-gene-CEs

The chip-seq data for histone modifications (H3K4Me1, H3K4Me2,H3K4Me3, H3K9ac, H3K9Me3, H3K27ac, H3K27Me3, H3K36Me3, H3K79Me3,H4K20Me1) were downloaded from UCSC (http://genome.ucsc.edu/encode/downloads.html) which is based on a direct submission from ENCODE production data (Human Genome Build 37 (hg19)). Specifically cell-line GM12878 data for histone modification board peaks were considered. GM12878 cell-line is mentioned as having a relatively normal karyotype. Also DNaseI hypersensitive site hotspot data for 2 replicates were downloaded from the above mentioned link.

Since the originally submitted ENCODE data was based on Grch37 human genome build, the coordinates needed to be upgraded to the latest version of human genome which is used in the analysis, which is Grch38. UCSC browser lift-over(https://genome.ucsc.edu/cgi-bin/hgLiftOver?hgsid=696327251_5kBwkhosCaqhOqgcpSUNVALQIBaf<u>) was used to convert the ENCODE data coordinates to the latest coordinates.</u>

The converted coordinates of histone modifications and DNaseI hypersensitive sites were checked for overlaps with the primate common CNSs and ncRNA. The median signal strength value was considered as the most reliable measure of the functional signature strength across CNSs and ncRNA-gene-CEs, since the distribution of the signal strength values were not normally distributed. Normality test for skewness and kurtosis was performed with shapiro-wilks test.

In order to determine the difference between conserved regions with respect to neutrally evolving regions in the genome,we also picked random samples of sequence coordinates from rest of the noncoding regions of human genome. We made 25 coordinate datasets for each histone modification. These random regions were picked on a very stringent criteria that they do not overlap with coding regions, already determined conserved noncoding regions or repeat regions of human genome. These random sequence sets contain same number of sequences, same length as the conserved noncoding sequences and also were retrieved from the same chromosome as a particular CNS.

### Selection pressure on primate common ncRNA-gene-CEs and CNSs

We downloaded hapmap human single nucleotide polymorphism data (ftp://ftp.ensembl.org/pub/release-94/variation/vcf/homo_sapiens/) for Yoruba population in Ibadan, Nigeria (YRI), Han Chinese in Beijing (CHB), China and individuals with Northern and Western European ancestry (CEU) collected from Utah. Then we determined how many SNPs overlap with primate common CNSs and primate common ncRNA-gene-CEs. Random coordinate sets from human genome were picked in order to compare between conserved regions and non-conserved regions. The random samples have the same number of instances, same length, and were picked from the same chromosome as the CNSs or the ncRNA gene. The random samples were normalized and tested for statistical significance by chi-square test.

### Evolutionary rates of CNSs and ncRNA-gene-CEs

The evolutionary rates for all primate common CNSs and ncRNA-gene-CEs at each branch were estimated by constructing neighbour-joining trees (Saitou and Nei. 1987) with 1000 bootstrap replications and evolutionary distances were calculated with maximum composite likelihood method with MEGA 5.0 (Tamura et al. 2011). The objective of this analysis was to determine if these CNSs and ncRNA-gene-CEs have gone through a fast-evolving phase before stabilizing and also which branches show accelerated evolution.

### Functional classification of genes in close proximity to CNSs and ncRNA-gene-CEs

First we determined the genes that are closest to CNSs and ncRNA-gene-CEs. These closest genes in the reference genome (Human) were considered as the likely target gene, if they had an orthologous gene in the most distal outgroup species used in the study (Mouse lemur). For these gene sets we determined functional enrichment via PANTHER14.1 classification system (Thomas et al. 2003).

### Likely target gene tissue expression patterns

In order to determine if the likely target gene tissue expression patterns follow the same trend as Gene Ontology (GO) analysis we retrieved Riken Fantom 5 project data for adult human and fetal data for 76 tissues through EBI (https://www.ebi.ac.uk/gxa/home) expression atlas. For this analysis we used 8089 likely target genes for CNSs and 7421likely target genes for ncRNA-gene-CEs that were found in the expression data. The CNSs and ncRNA in the genome function differently, the CNSs mostly function as cis regulatory elements whereas ncRNA is known to function as cis and trans elements. The objective of this analysis was to determine if cis and presumably trans acting elements are close in proximity to similar types of genes. The genes that had any sign of expression were considered for further analyses. The number of genes expressed for brain and nervous system related functions and several tissues related to housekeeping functions were considered as two potential groups for better comparison. Also we constructed 10 random gene sets for each group (CNSs and ncRNA-gene-CEs) for clarity on random expectation. The statistical significance of the data and random samples were determined by one-sample t-test with statistical significance level of 0.05.

### Clustering patterns of CNSs and ncRNA-gene-CEs

We determined the tendency of primate common CNSs and conserved ncRNA occurring in clusters. This analysis was conducted by considering groups of CNSs and ncRNA-gene-CEs in clusters with one likely target gene (considering closest orthologous gene as described before). In other words this analysis determines how many CNSs or conserved ncRNAs are associated with its closest orthologous target gene and their occurrence trends in the reference genome. We classified the CNSs and ncRNA-gene-CEs into two groups for further comparisons, that is genes with 1-2 CNSs / 1-2 ncRNA-gene-CEs and genes with >6 CNSs / >6 ncRNA-gene-CEs.

### Tissue expression levels for genes associated with CNSs and ncRNA-gene-CE clusters

Furthermore we determined if genes with many CNSs or ncRNA have difference in their expression level in diverse brain related tissues. Here we used genes expressed in four tissues namely amygdala, caudate nucleus, cerebellum and spinal cord. To have an elaborate view on tissue expression we considered genes that showed any kind of expression without a baseline TPM (Transcripts per Million) threshold. The objective of this analysis is to see any differences or similarities in gene expression patterns for CNSs and conserved ncRNA in clusters and how the 2 groups are functioning on the genomic level. This helps to shed light upon understanding the evolutionary dynamics of CNSs and conserved ncRNA with respect to expression. Statistical tests for significance were determined by Mann-Whitney U test.

### CNSs and ncRNA-gene-CEs relation to experimentally verified enhancers, promoters and CTCF binding sites

We determined if our primate common CNSs and ncRNA-gene-CEs are actually related to predicted enhancer, promoter and CTCF binding sites in (ensemble.org/info/genomeensembl.org/info/genome/funcgen/regulatory_build. html). Based on premise, if CNSs are actually governing gene regulation we expect more enhancers to be associated with CNSs and also checking the magnitude of ncRNA-gene-CEs association to enhancers helps to elucidate the functional landscape of conserved ncRNA. Whereas CTCF binding sites are concerned, they are considered important and conserved across lineages functioning as insulator elements, blocking enhancer activity functioning as domain barriers (Hou et al. 2008; Cuddapath et al. 2009;Herold et al. 2012).

Promoters help to get an understanding on which regions have higher tendency to get transcribed, by comparing association of our primate common CNSs and ncRNA-gene-CEs to documented promoter regions help to differentiate CNSs and ncRNA-gene-CEs and their specific role in a genome.

We compare our dataset against random regions with same length and location (chromosome) as CNSs or ncRNA-gene-CEs of the reference genome for better clarification on how well we can be confident with the functionality of these conserved regions. Statistical significance against random regions were determined by one sample t-test.

Also to be confident that the pattern of representation is not dependent on the initial number difference in CNSs and ncRNA-gene-CEs, we further randomly picked 20 samples of 59,870 CNSs (same number as identified ncRNA-gene-CEs) and searched for the signals of enhancers, CTCF binding and promoters in the 20 sampled datasets. Also we determined the functional categories for the likely target genes of CTCF binding, enhancer and promoter overlapping CNSs and ncRNA-gene-CEs.

### Determining the presence of primate deep conserved elements in mammals, i.e more ancient conservation

We tried to determine how many of the primate common conserved elements are found in mammalian common ancestor. For this analysis we searched primate common CNSs and ncRNA-gene-CEs separately in rat (*Rattus norvegicus*) and mouse genomes (*Mus musculus*) in Ensembl release 94. The likely target gene GO was determined via PANTHER14.1. Also we looked at the tissue expression patterns of these genes.

### Have Hominoid specific conserved non coding elements originated to render specific functions to hominoid lineage?

We determined the Hominoid lineage specific elements that are unique to human, gorilla, orangutan and gibbon, by searching for homologs of this group in all primate out-group species. Any sequence instance that was found in any of the out-group species were removed from the final data set or any further analyses. Finally we considered only the elements above 90% percentage identity for further analyses. These hominoid unique elements were checked for their closest likely target gene, related functions and tissue expression patterns.

### Do CNSs and ncRNA conserved elements belong to independent categories?

Since we observed similar functional categories for CNSs and ncRNA conserved elements we identified in the study, it was important to know if there were any CNSs, ncRNA cross homology sequences causing this scenario. Therefore we searched all the conserved noncoding elements against itself with Blastn with e<0.001. After removing self-hits and one copy of reciprocal hits the remaining hits with a CNS and ncRNA combinations were extracted (4772 sequence instances). The coordinates of these CNS, ncRNA cross homology entities were searched for their features with Ensembl feature category based on annotation.

## Results

### Identification and characterisation of a highly conserved CNS and ncRNA dataset spanning primates

To generate a conservative, high confidence set of CNS and ncRNA sequences, we screened for conserved noncoding regions present in 12primates. This yielded 153,475 primate common conserved noncoding regions. Of these, 59,870 are candidate ncRNA gene overlaps (overlapping with annotated ncRNAs) while 93,605 elements were found in other non-coding (Intergenic, Untraslated Regions [UTR], introns of protein coding genes) regions of the genome. Based on these assignments, we will refer to these noncoding regions as ncRNA -gene-CEs and CNSs, respectively.

### About 40% of primate common elements are found in Mammalian common ancestor

We found 35972, CNSs and 23549 ncRNA-gene-CEs that are common to the 12 species that we have used in the study (Figure 7 A). This means only 38% of CNSs and 39% (Figure 7B) of the ncRNA-gene-CEs found as primate common elements already existed in the mammalian common ancestor, which are very ancient elements. Therefore majority (about 60%) of primate common elements we identified in this study appear to have solely originated in the primate common ancestor.

The closest likely target gene functional classification for these elements revealed that mammalian common ncRNA-gene-CEs (while assuming a cis activity for these elements) are related to regulation of synapse structure and activity (Supplementary Table 2). Mammalian common CNSs too show similar pattern of being related to central nervous system but with less fold-enrichment meaning that these ancient noncoding conservation is not the sole determinant of brain function in primates. i.e primate evolution has taken a new turn by recruiting or generating new regulatory elements to suits its lineage necessities.

### Purifying selection on conserved noncoding regions

In order to test for the level of selection of primate common, CNSs and ncRNA-gene-CEs we carried out the derived allele frequency analysis. Functional regions under purifying selection are expected to have lower levels or lower frequency of derived alleles than regions that are neutrally evolving. Our results show that the primate common CNSs and ncRNA-gene-CEs both have a high percentage of lower derived alleles than neutrally evolving random sequences. Alleles with lower frequency levels (DAF < 0.1) in both CNSs and ncRNA-gene-CEs show significantly higher level of abundance than expected. (Z-test p<0.00001). Also it is important to note that high frequency alleles are less abundant in evolutionarily conserved regions than neutrally evolving background sequences of the human genome. Abundance levels of SNPs with derived allele frequency>0.9 are significantly higher (z-test p<0.00001) in neutrally evolving regions compared to conserved regions in our results. Also it is clear that ncRNA-gene-CEs are under slightly higher levels of purifying selection than CNSs. This scenario is uniform with both Yoruba and Han Chinese population (Supplementary Figure 1). With Yoruba population, CNSs had a percentage level of 47.65% (Figure 4A) for lower allele frequency whereas ncRNA-gene-CEs had 49.81% (Figure 4B). Han Chinese and European (Supplementary Figure 1; Supplementary Figure 2) SNP data also showed that ncRNA had a higher percentage level (24.3%, 50.02%) compared to CNSs for lower derived alleles (<0.1). For Yoruba population the statistical significance levels of CNSs and ncRNA-gene-CEs SNP distribution from the random expectation are p < 2.05854E-49, p <8.91508E-74(chi-square test) respectively. In other words the observed SNP distribution in CNSs and ncRNA-gene-CEs has a very low probability of being a chance distribution.

**Figure 3:**
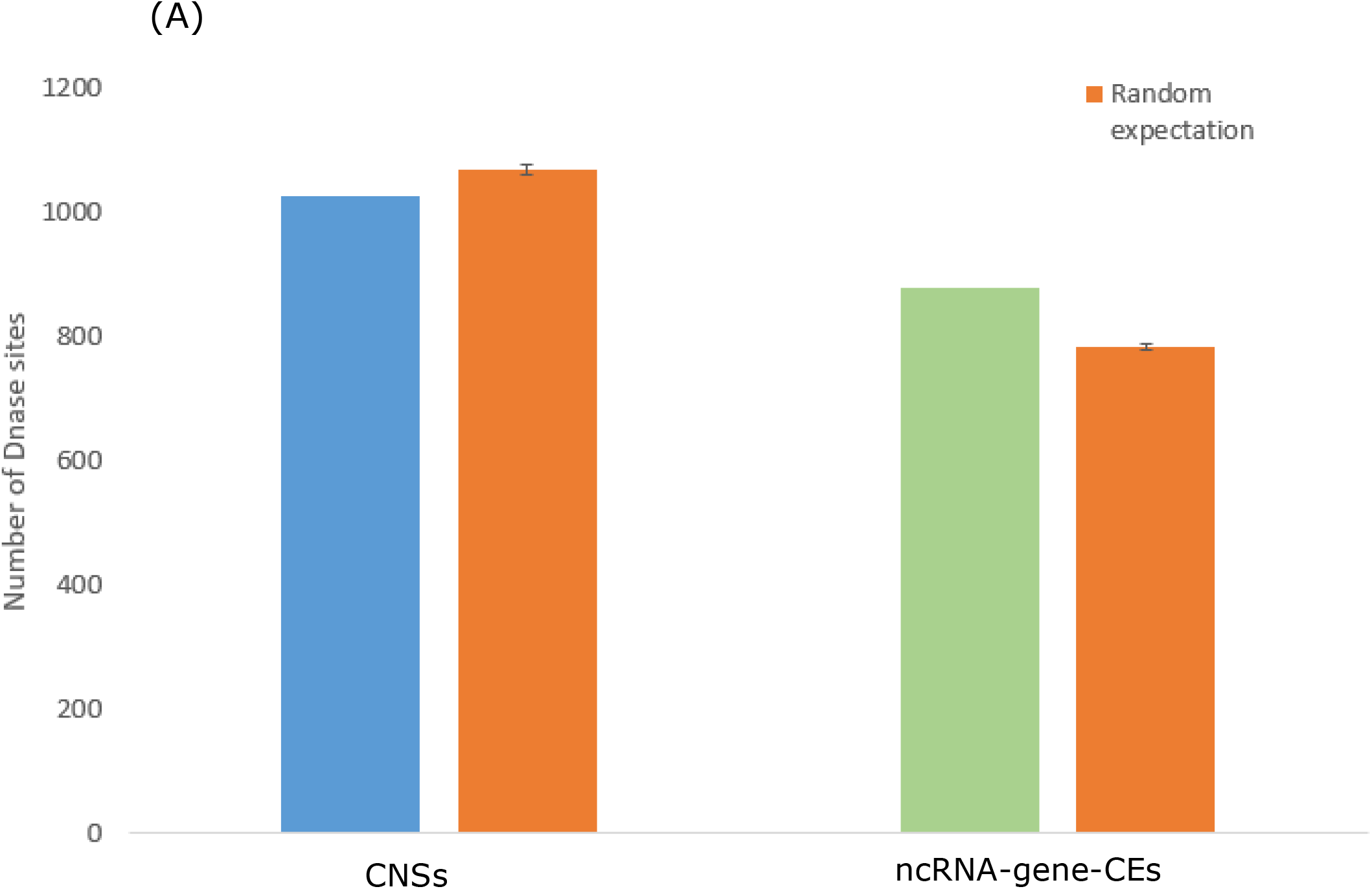
Dnase I hypersensitive sites associated with primate common CNSs and ncRNA-gene-CEs. The sites that are found inside conserved regions were considered. The random expectation was determined by 25 random samples from human genome that are not conserved and are presumably neutrally evolving. The results were statistically significant (p<0.0001).

**Figure 4:**
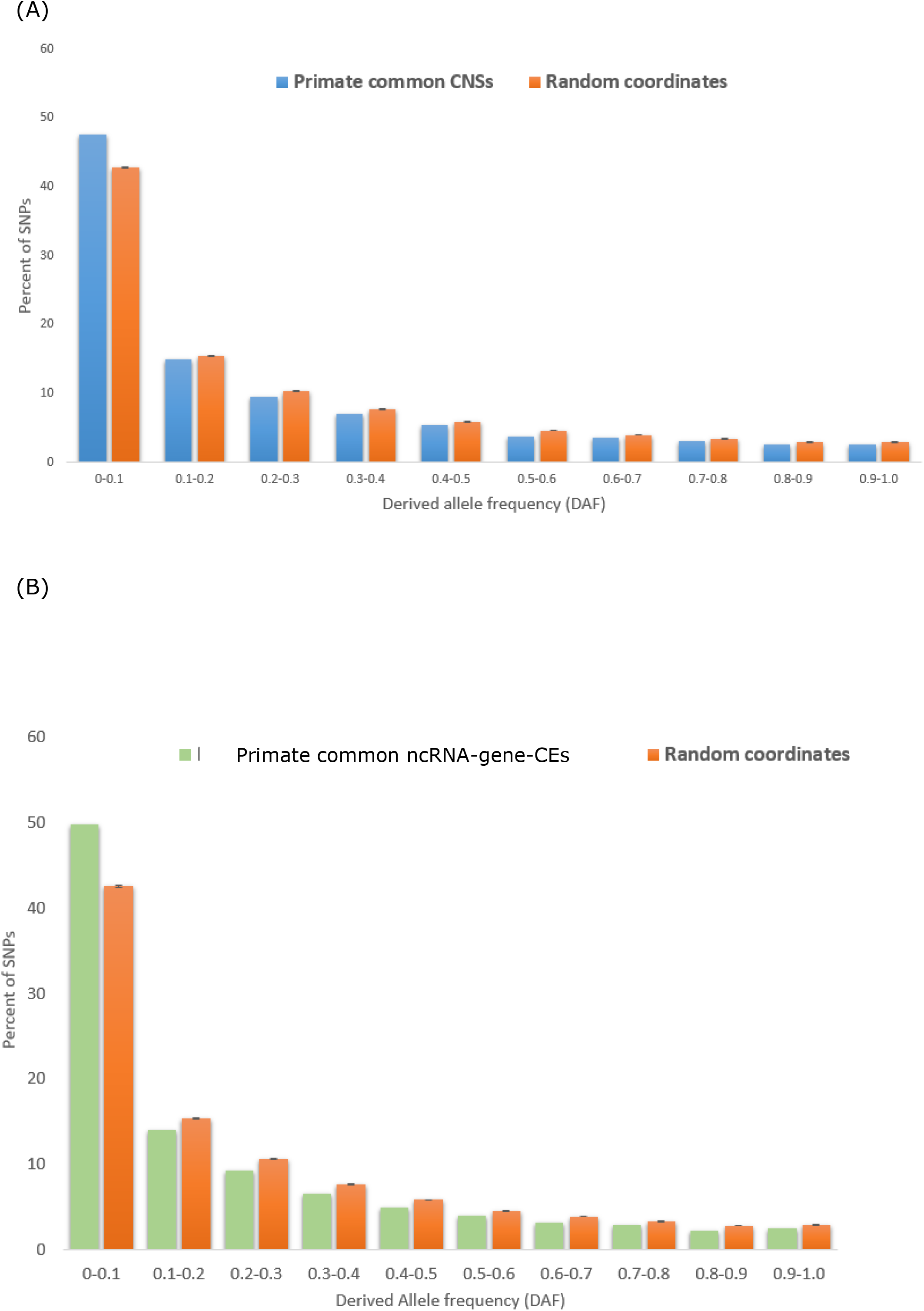
Derived allele frequency analysis for primate common CNSs and primate common ncRNA-gene-CEs using Yoruba population. Random coordinates from the human genome build Grch38 were picked and considered as neutrally evolving regions compared to conserved regions. The Hapmap project SNP data was used for this analysis. CNSs and conserved non-coding RNA both have higher percentage of SNPs for lower derived allele frequency level <0.1 compared to random expectation. The results are statically significant at p <0.05 (z-test).

These results show that generally, conserved non-coding regions are under purifying selection and some conserved non-coding regions, such as primate common ncRNA-gene-CEs in this analysis are under a slightly higher selective pressure than CNSs. It is already well known that conserved non-coding regions are functionally important and disruption or mutations of these conserved regions can lead to detrimental effects.

Primate common ncRNA-gene-CEs also show a higher level of selection or in other words, less tolerant of higher frequency mutations. Majority of the primate common ncRNA-gene-CEs we identified in this analysis were long non-coding RNA. It has been shown in many studies that long non-coding RNA can also function in gene regulation controlling gene expression (Bao et al. 2013), therefore it can be expected that conserved ncRNA to be under purifying selection.

### Functional classification of genes associated with CNSs and ncRNA-gene-CEs

Functional classification for the closest gene for CNSs showed a pattern where majority of the genes were associated with neuron development (Table 1A). Surprisingly the closest gene to ncRNA-gene-CEs also followed the same pattern (Table 1B) in gene ontology. The genes closest to ncRNA-gene-CEs were also involved in neuron development and brain related functions. Even though the fold enrichment (genes observed, over expected number) was slightly higher for genes closest to CNSs compared to ncRNA-gene-CEs, this uniform result signifies that these ncRNA-CEs may be involved in cis regulation of these genes. It definitely is intriguing that these primate common CNSs and ncRNA-gene-CEs follow similar patterns in their potential functions.

**Table 1.** – GO terms related with closest genes for CNSs and ncRNA-gene-CEs. (A) Gene Enrichment values for likely target genes for CNSs (B) Gene Enrichment values for likely target genes forncRNA-gene-CEs

| **GO term** | **Fold enrichment** | **P-value** |
| --- | --- | --- |
| Regulation of cell differentiation | 1.72 | 1.13E-03 |
| Neuron differentiation | 1.71 | 3.43E-07 |
| Regulation of developmental processes | 1.70 | 1.97E-04 |
| Cell morphogenesis involved in neuron differentiation | 1.70 | 5.20E-04 |
| Nervous system development | 1.62 | 5.59E-08 |

**(B)**
| **GO term** | **Fold enrichment** | **P-value** |
| --- | --- | --- |
| Positive regulation of transcription by RNA polymerase II | 1.58 | 1.04E-05 |
| Neuron differentiation | 1.56 | 7.16E-05 |
| Generation of neurons | 1.54 | 5.88E-05 |
| Neurogenesis | 1.52 | 9.06E-05 |
| Nervous system development | 1.47 | 5.16E-05 |

### Likely target gene tissue expression patterns for primate common CNSs

Majority of the closest proximity genes for both CNSs and ncRNA-gene-CEs were related to brain and nervous system related tissues. Amygdala, brain meninx, occipital lobe and spinal cord among others showed the highest fraction of expressed genes for both categories, CNSs and ncRNA-gene-CEs (Figure 6A; 6B).The expressed gene fractions are statistically significant in all cases compared to random samples (p<0.00001).This pattern is same as what was observed in the GO analysis where high level of enrichment was shown for GO term nervous system development. Also random gene sets that are not closest to CNSs or ncRNA-gene-CEs, showed significantly lower levels of genes expressed in brain related tissues compared to closest likely target genes. Also there is a significant difference in CNS and ncRNA percentage gene expression for brain related tissues, that is CNSs have a slightly higher tendency to harbour brain or nervous system related genes as the adjacent likely target gene compared to ncRNA-gene-CEs(P = 0.00799, one tail t-test)

### Hominoidae specific CNSs and ncRNA-gene-CEs are less associated with immune responses but more with neuron development

We identified a total of 12481 elements that presumably originated and are conserved only in the hominoidae lineage. Of the total elements 3363 were ncRNA-gene-CEs, while 9118 were CNSs. In general terms these lineage specific elements also cluster close to genes that are predominantly related with neuron development and differentiation (Supplementary Table 3).

### Accelerated evolution of conserved noncoding and ncRNA-gene-CEs

We found primate common CNSs and ncRNA-gene-CEs have gone through a phase of accelerated evolution in several different stages during evolution before getting stabilized. The human-gorilla common ancestor sequences showed 4 times faster evolutionary rate compared to human-gorilla-orangutan common ancestor for both CNSs and ncRNA-gene-CEs (Figure 5A, 5B). Also we observed that Hominoidae common ancestor showed accelerated evolution in CNS sequences after diverging from Hominoidae-old world monkey common ancestor. Also old world monkeys show about 2 fold acceleration after diverging from Hominoidae-old world monkey common ancestor.

**Figure 5:**
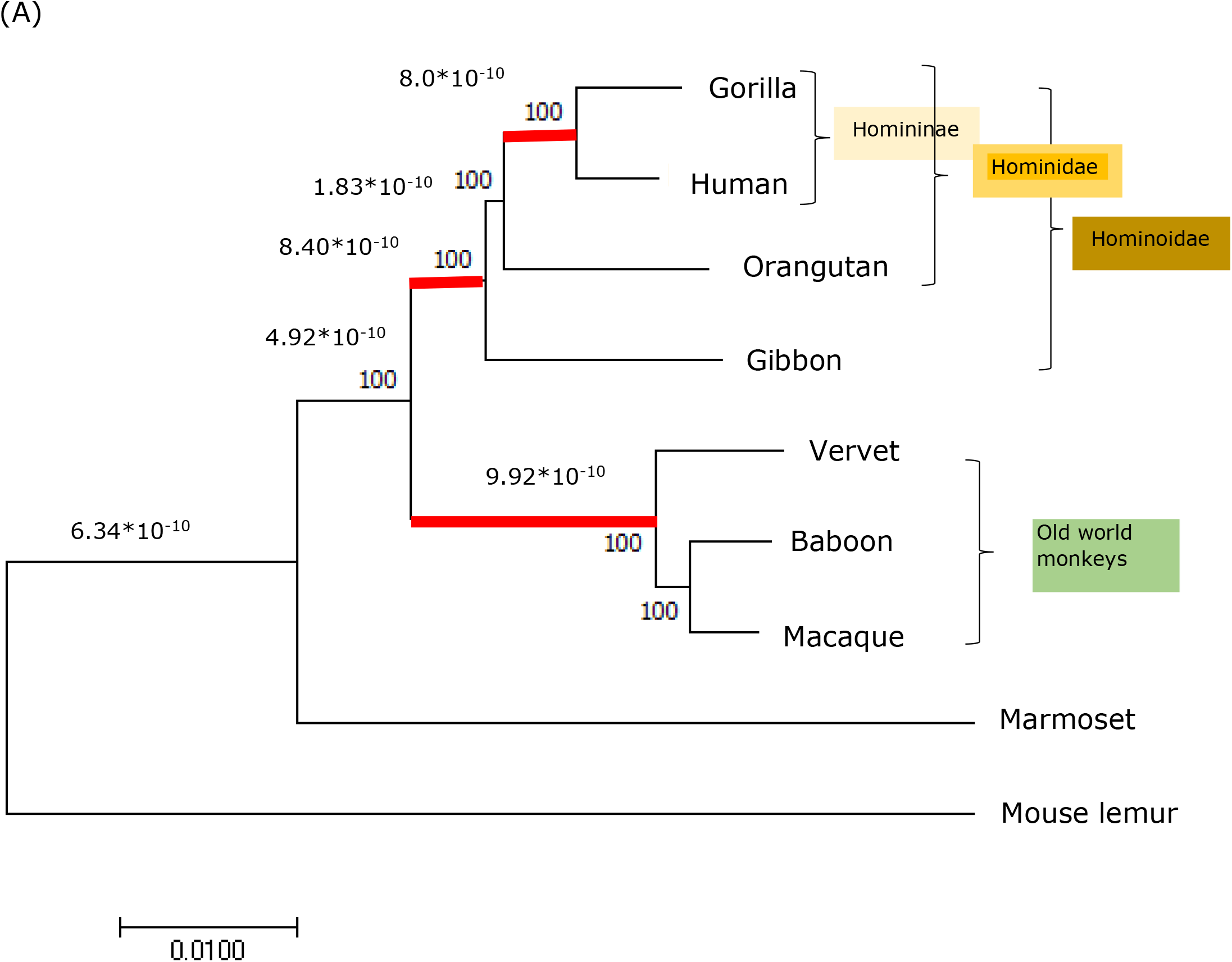
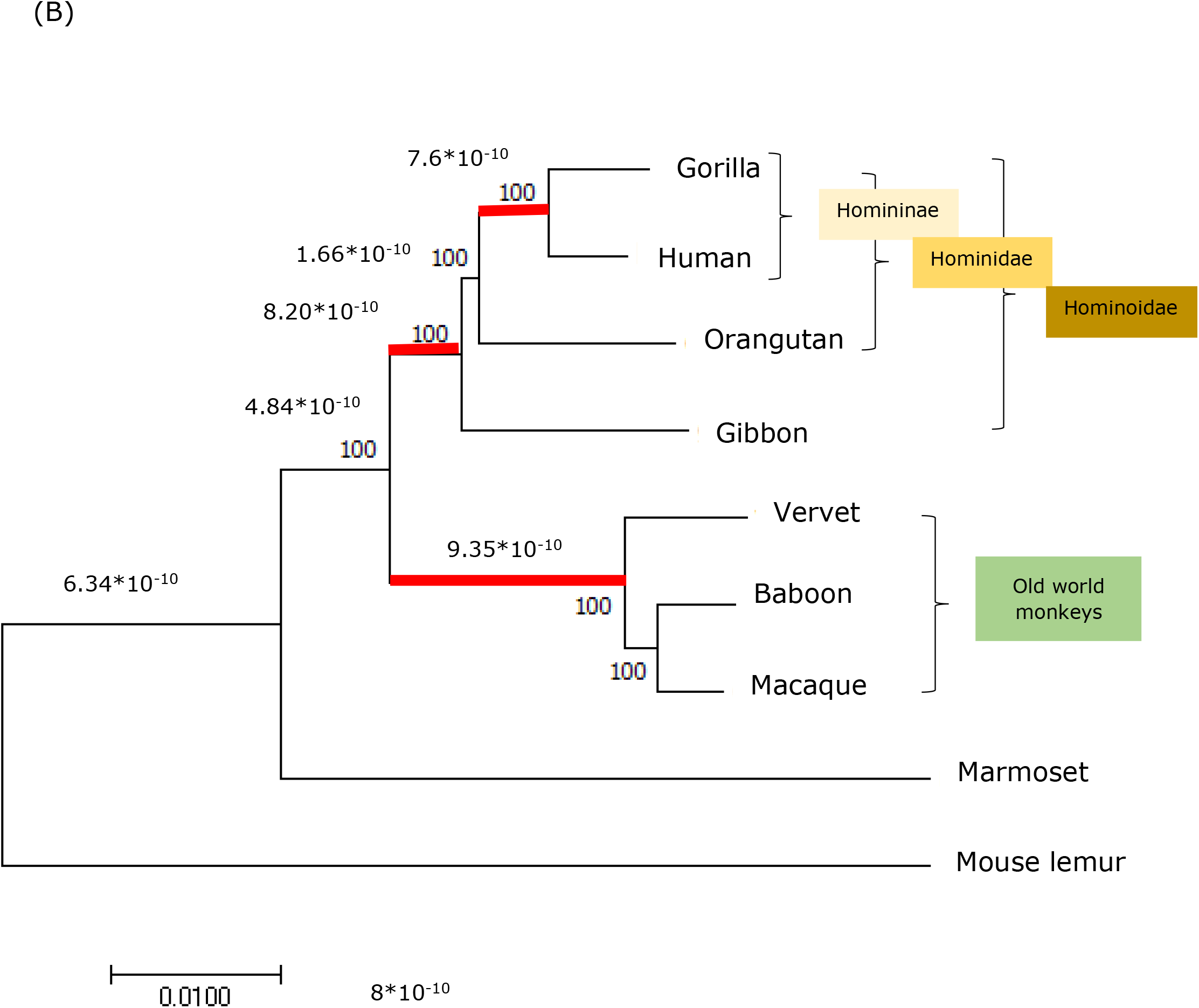
(A) CNSs show faster evolution at different stages in phylogeny. (B)Evolutionary rates of ncRNA-gene-CEs in primate lineage. The branches with faster substitution (>=8*10^-10^per site per year) are highlighted by a red bar. Primate common CNSs went through acceleration at Human-Gorilla, Hominoidae and Old world monkey common ancestor.Conserved ncRNA gene sequences also show acceleration during primate evolution. The mutation rate is as per site per year.

**Figure 6:**
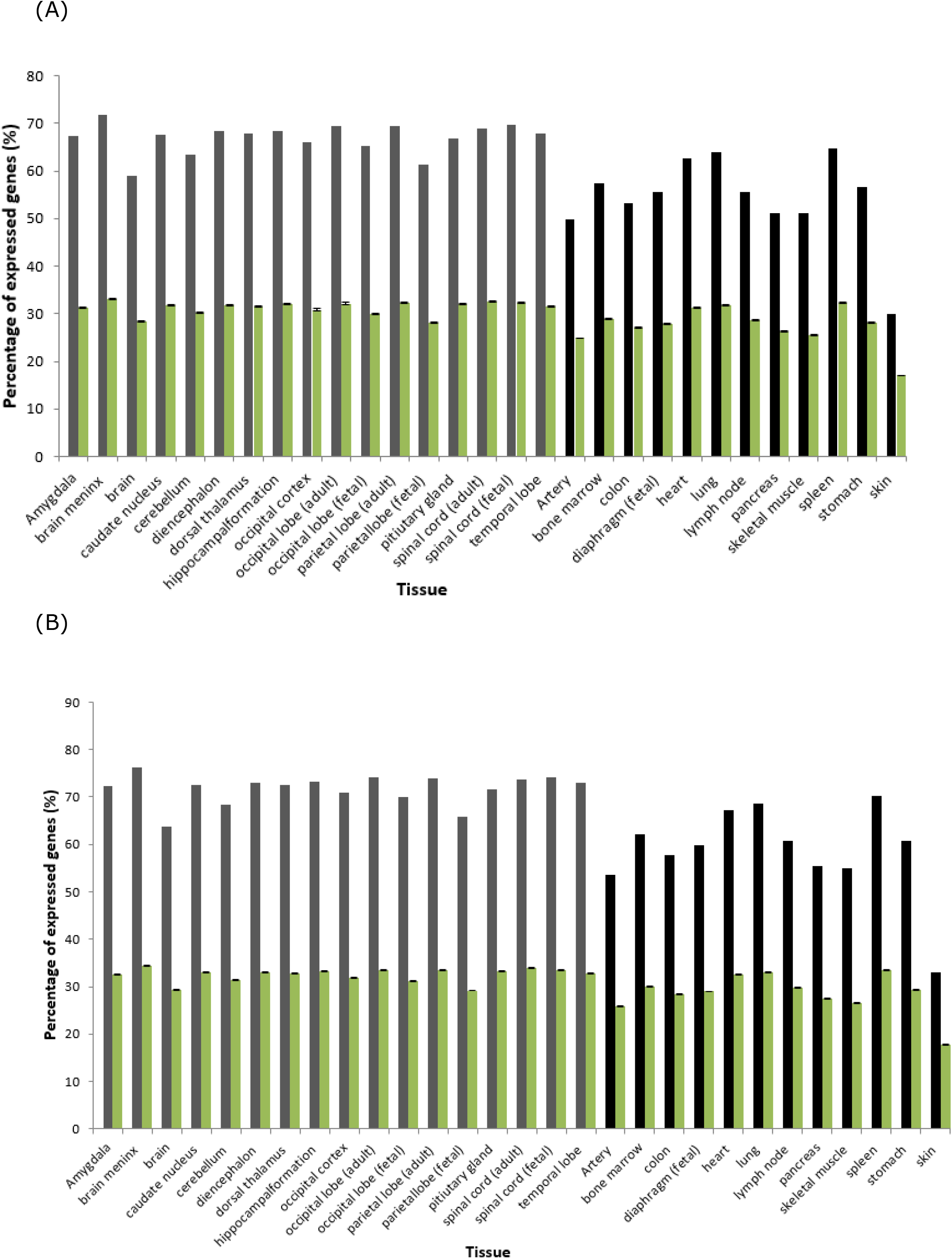
Primate common CNSs and ncRNA-gene-CEs closest gene expression patterns. (A) The percentage of expressed genes for CNS. (B) The percentage of expressed genes for ncRNA-gene-CEs. Grey, black bars represent brain related and house-keeping tissues respectively. Green bars signify the random expectation. Statistical significance of all results is p<0.00001.

**Figure 7:**
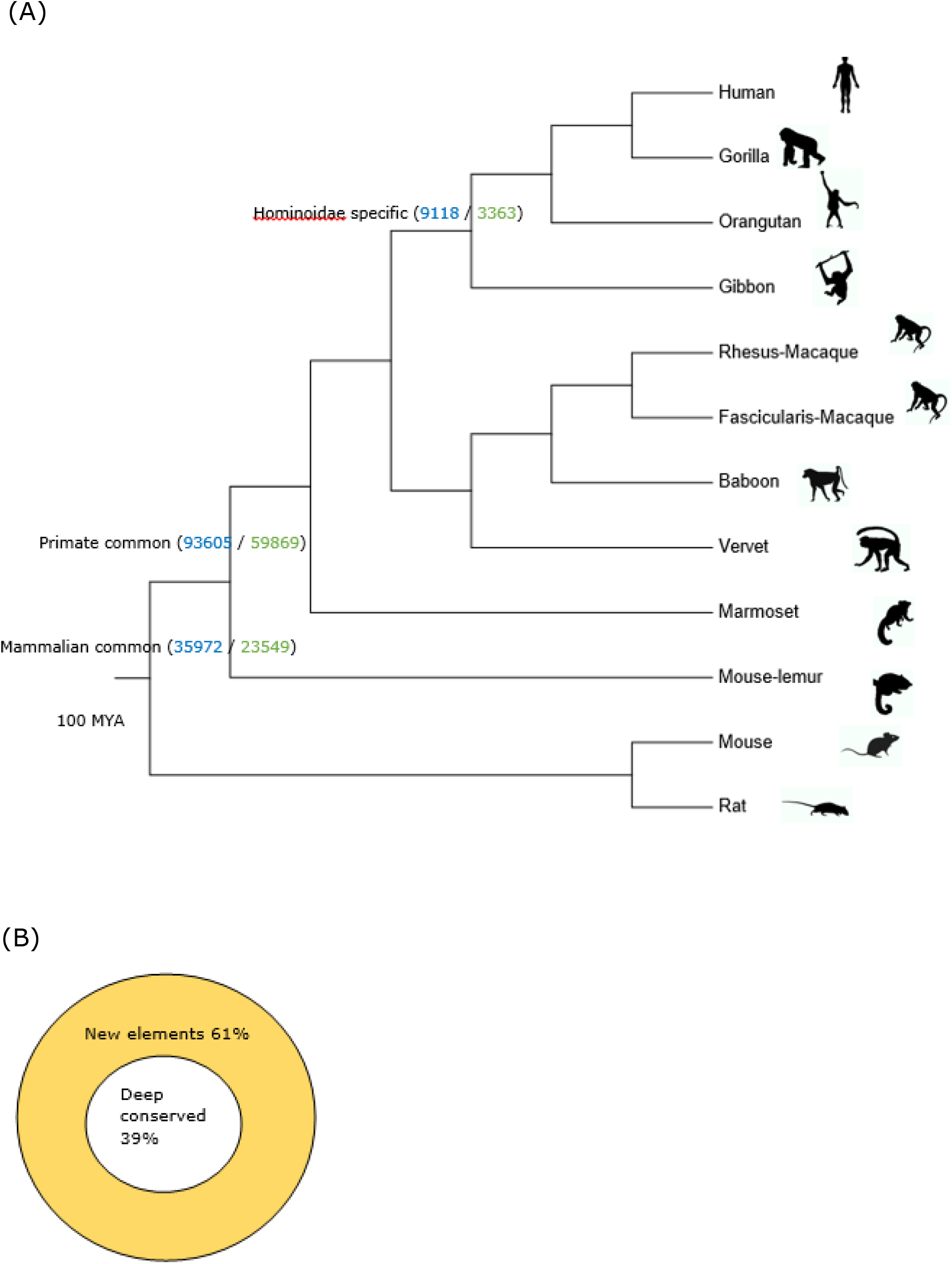
(A) Number of CNSs and ncRNA-gene-CEs found in primate, mammal and hominoidae common ancestor. (B) percentage of primate common newly originated elements and deep conserved elements. The numbers in blue on respective branches represent CNSs and ncRNA in green. MYA-Million Years Ago

### Epigenetic profiles of conserved primate CNS and ncRNA genes

H3K4Me1 is a histone modification associated with active enhancer regions (Koenecke et al. 2016; Barakat et al. 2018). The median signal strength of H3K4Me1 located inside CNSs were slightly higher but not statistically significant compared to random coordinates (Figure 2A). Also another histone modification that is associated with activated enhancer regions is H3K27ac. The CNSs have significantly higher levels of H3K27ac compared to random samples (p < 0.0001). H3K79 methylation is known to be associated with developmentally regulated gene expression in multicellular eukaryotes. For this histone modification, both primate common CNSs and ncRNA-gene-CEs have high levels of signal strength compared to random samples. H3K9ac which is a histone modification mark predominantly found in active promoter regions showed high signal strength in CNSs with respect to random sample (p = 0.003). In general according to our results, CNSs show a higher tendency to be active enhancer elements associated with regulatory activities.

H3K4Me1 in ncRNA-gene-CEs were statistically significantly lower than random expectation (p < 0.0001), signifying that these primate common ncRNA-gene-CEs may have less chance of functioning as active enhancers. Also H3K27ac, another active enhancer histone modification showed significantly less strength in ncRNA-gene-CEs compared to random samples. This result implies that the ncRNA-gene-CEs in this case may not be or less associated with active enhancers as CNSs.

One interesting result is that H4K20Me1 show significantly higher levels of association to primate ncRNA-gene-CEs compared to CNSs (Figure 2B). This histone modification is involved in DNA replication and damage repair. *CCND1* is the first long ncRNA that was identified as being transcribed in response to DNA damage signals (Wang et al. 2008; Klein and Assoian. 2008). *LincROR* has been identified as a P53 repressor in response to DNA damage (Zhang et al. 2013). In addition several other studies have revealed ncRNA association to DNA repair and replication (Ge and Lin. 2014; Hawley et al. 2017). The fact that we found that primate common ncRNA-gene-CEs had a high signal strength for H4K20Me1 in a way further signifies association of ncRNA-gene-CEs with DNA damage repair.

The number of histone modification sites found inside CNSs and ncRNA-gene-CEs are given in Supplementary Table 1.

### DNaseI hypertensive sites (DHSs) in conserved noncoding regions

We found that CNSs have significantly less abundance of DNaseI hypersensitive sites associated with them (p<0.0001), compared to random expectation. Whereas conserved ncRNA show an opposite trend where they have more DNaseI sites in them (p<0.0001), compared to random samples (Figure 3). DHSs signify open chromatin regions according to many reports (Song et al. 2011; Madrigal and Krajewski. 2012; Li and Cui. 2018). Therefore this result implies that the primate common CNSs we identified tend to be located in regions with less possibility of being open chromatin compared to random regions in the genome. But primate common ncRNA-gene-CEs have more DHSs compared to the random expectation implying a higher tendency for these regions to be in open chromatin conformation.

### CNSs and ncRNA-gene-CEs both show less enrichment in CTCF binding and higher enrichment in enhancer regions compared to random regions in the genome

We see statistically significantly higher levels of enhancers in CNSs and ncRNA-gene-CEs compared to random expectation (p<0.05) (supplementary figure 3A). This signifies and further establishes the fact that in this case primate common CNSs and ncRNA-gene-CEs play a significant role as enhancer elements in these genomes. Interestingly we find that both CNSs and ncRNA-gene-CEs show less enrichment in CTCF binding compared to other random regions in the human genome (p<0.05), signifying these identified conserved regions have a less tendency to be insulator or repressive domains (supplementary figure 3B). But one important feature to note is that more promoter regions are associated with ncRNA-gene-CEs in our dataset compared to CNSs, which reflects more transcriptional activity (p<0.05) of ncRNA-gene-CEs compared to CNSs (supplementary figure 3C).

In comparison the randomly picked (same number of CNSs as ncRNA-gene-CEs in the original data set – sampled 20 times with replacement) CNSs with ncRNA-gene-CEs, showed that number of CNSs that harboured enhancer were not statistically significant compared to ncRNA-gene-CEs, meaning that these ncRNA-gene-CEs and CNSs have equal tendency to have enhancer elements (supplementary figure 3A). Also Sampled CNSs showed no statistical significance in abundance of CTCF binding sites compared to ncRNA-gene-CEs, signifying that CNSs and ncRNA-gene-CEs have equally less tendency to function as insulator elements (supplementary figure 3B). But promoter regions were significantly less abundant in CNSs than ncRNA-gene-CEs signifying a difference in transcription activity in CNSs and ncRNA-gene-CEs (p is < .00001) (supplementary figure 3C) which also agrees with the comparison we made to randomly picked non conserved regions of the genome to primate common CNSs and ncRNA-gene-CEs. Also the enhancer and CTCF overlapping CNSs showed higher enrichment related to nervous system and brain related structural and functional GO terms. But promoter overlapping CNSs and ncRNA-gene-CEs were more related with housekeeping structural and functional terms (supplementary table 6). This further clarifies experimentally verified enhancer related CNSs are related to brain and nervous system function as we have noted earlier computationally.

### CNSs and ncRNA-gene-CEs are mostly mutually exclusive groups of conserved elements

We found that 4772 conserved elements showed CNS-ncRNA cross homology, In other words only 3.10% of all the identified conserved non coding elements fell into this category. The feature categories of these cross homology elements were predominantly categorized as miRNA target sites (Figure 8, Supplementary table 5). Also it’s important to note that miRNA target site coordinates are all non-overlapping and are located in diverse locations in the reference genome. This result clarifies that the CNS, ncRNA conserved elements predominantly fall into mutually exclusive groups. This further clarifies that the previously observed Similar functional categorization of the CNS and ncRNA conserved elements could not have been a result of cross homology elements that is only a small fraction of the dataset.

**Figure 8:**
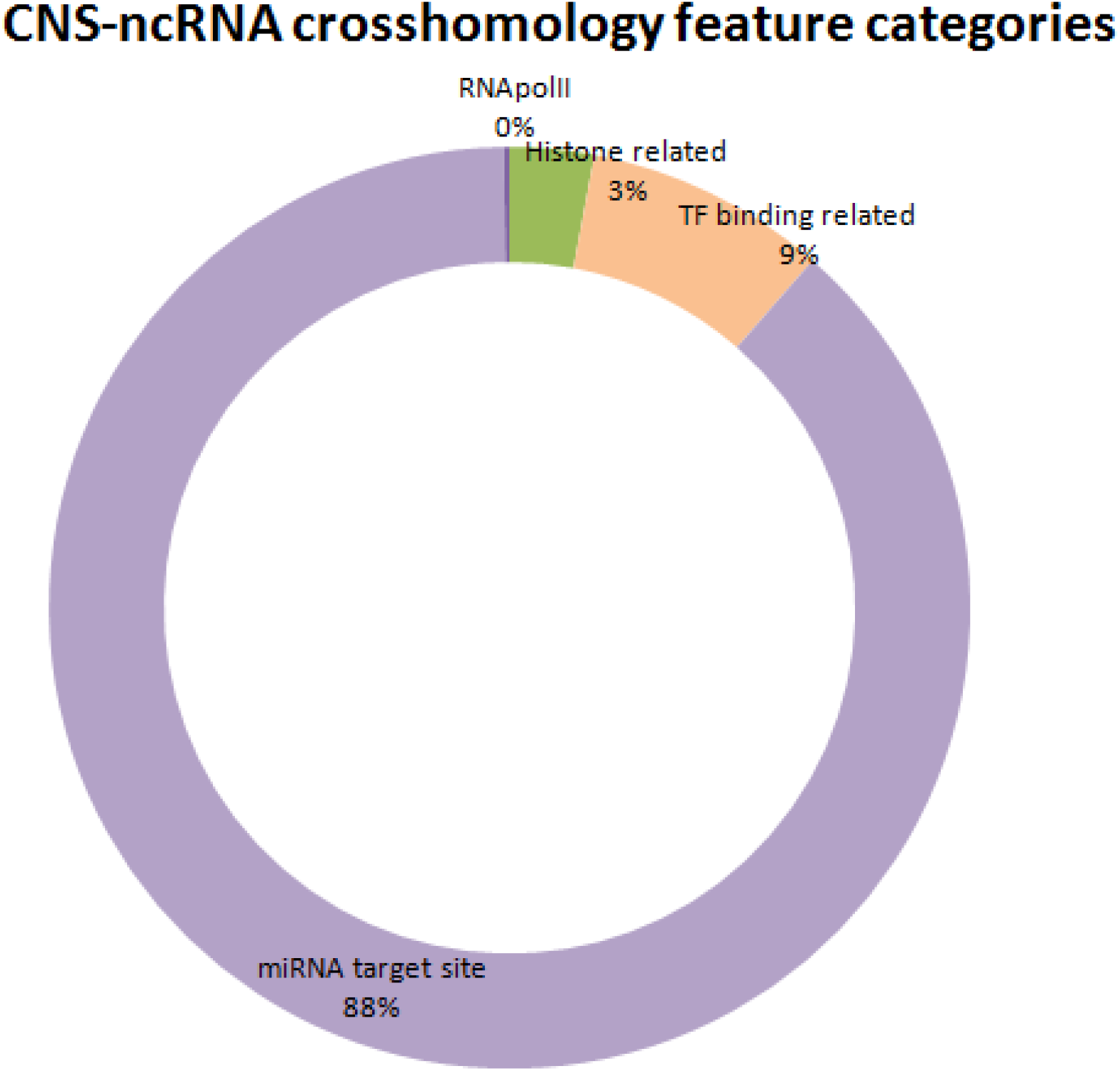
Feature categories based on Emsembl annotation for the CNs-ncRNA 4772 cross homology elements. Only 3.10% of all elements fall into cross homology category. Majority of the elements that show cross homology are miRNA target sites.

### Clustering patterns of CNSs and ncRNA-gene-CEs

We found that majority of the cases were associated with one conserved non-coding region with one likely target gene, for both CNSs and ncRNA. But ncRNA had slight higher (27%) of its likely target genes being associated with only one conserved ncRNA (supplementary figure 4). Never the less CNSs and conserved ncRNA showed no statistical significance regards to occurrence in clustering, implying that it is equally likely for CNSs and conserved ncRNA to occur in clusters.

### Gene Expression levels of clustered CNSs and conserved ncRNA reveals significant difference based on clustering pattern

We found 3380, 3457 likely target genes in total, associated with 1-2 CNSs and >6 CNSs respectively. 1-2 CNS group showed more genes being expressed in amygdala and spinal cord (Figure 9). In general 1-2 CNSs group showed more genes being expressed compared to genes associated with >6 CNSs. This implies that genes with more CNSs do not necessarily have a tendency to be expressed more often. Also we closely observed the expression levels of these genes and found that there is a significant difference in the gene expression levels for each of the 4 tissues for the 2 groups. More expression levels were seen for genes associated with 1-2 CNSs for all for tissues (amygdala, caudate nucleus, cerebellum and spinal cord) compared to CNSs in clusters of more than 6 conserved regions (Figure 9 A).

**Figure 9:**
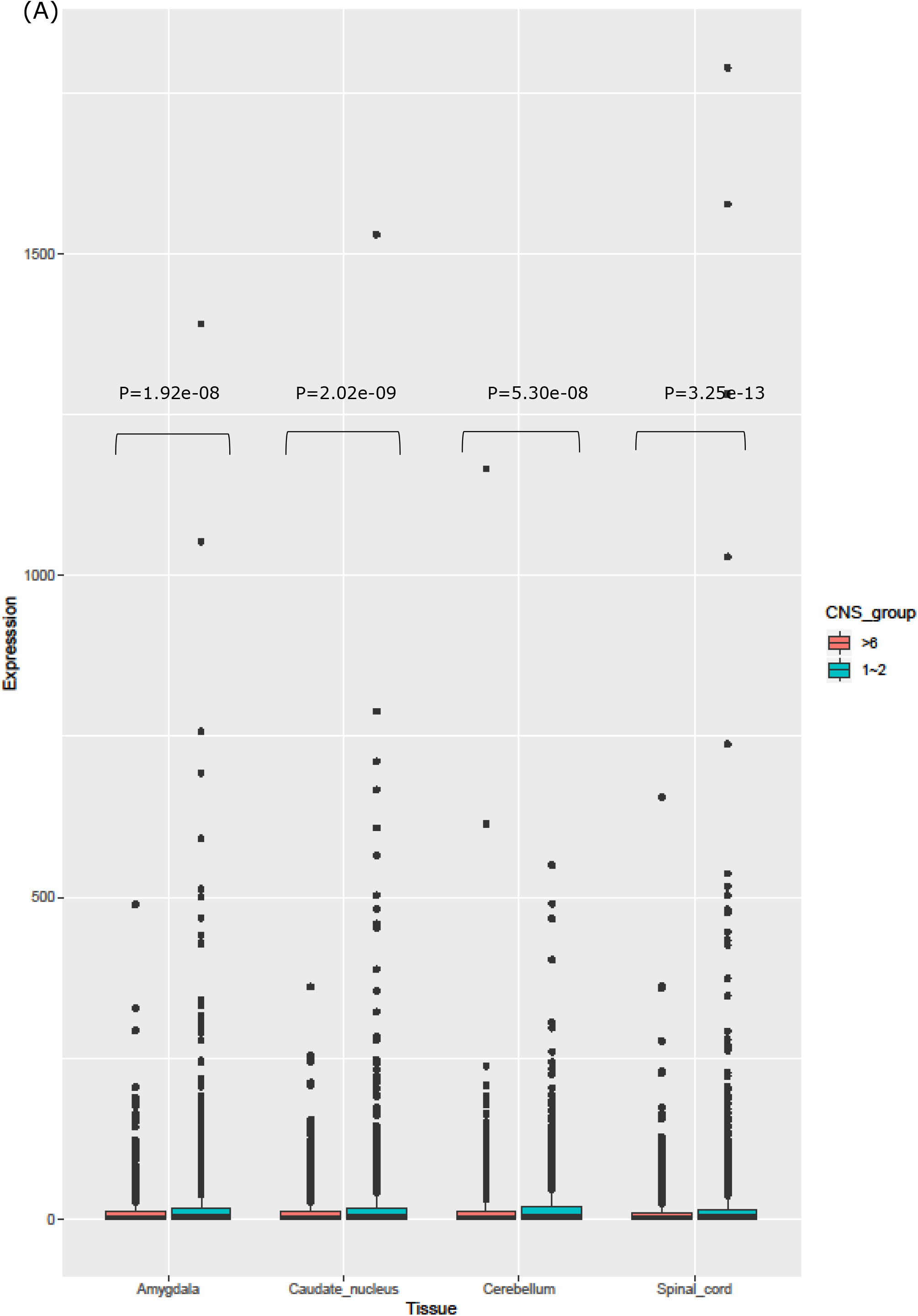
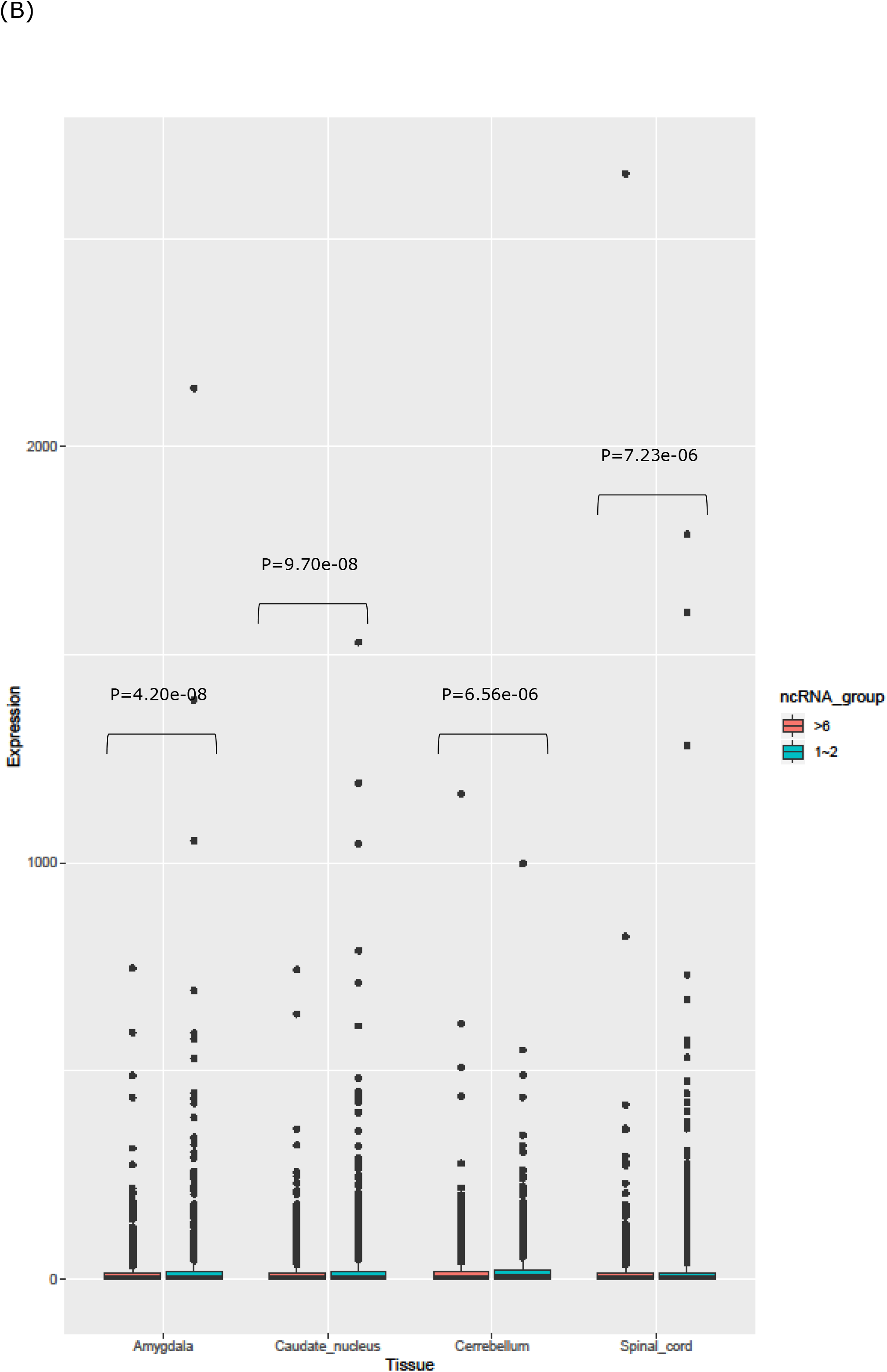
(A) Expression levels for genes associated with clustered CNSs.The CNS_group represents grouping pattern that was considered, i.e 1-2 means genes with 1or 2 CNSs, whereas >6 means genes with more than 6 CNSs. The statistical significance was determined by Mann-whitney U-test. The x-axis represents gene expression level for each of these groups for several brain related tissues (absolute values are used in TPM (Transcripts per Million)). **(B) Expression levels for genes associated with clustered conserved ncRNA.** The ncRNA group represents grouping pattern that was considered,i.e 1-2 means genes with 1or 2 conserved ncRNAs, whereas >6 means, genes with more than 6 conserved ncRNAs. The x-axis represents gene expression level for each of these groups for several brain related tissues.

Conserved ncRNA also followed the same pattern as CNSs where 1-2 conserved ncRNA cluster related genes showed higher expression levels compared to genes with clusters of more than 6 conserved ncRNAs (Figure 9B). It appears that multiple blocks of conserved ncRNA or CNS regions are not necessarily required to achieve high expression levels of the target genes.

Also further investigation showed that that the genes with 1-2 CNSs or ncRNA clusters were very much similar in their expression levels across other tissues we considered for this analysis.

## Discussion

We have identified a set of highly conserved ncRNAs and CNSs from primates. Many are also found in other mammals, and our analyses indicate these are under selective constraint, giving us confidence that they are likely to be functional elements.

We assessed the probable function of our CNS and ncRNA dataset by looking at the function of the closest adjacent gene. This is frequently done for CNSs, on the basis that their mode of action is local, and in cis (Nelson and Wardle 2013, Vavouri et al.2007), while we expected no clear signal for candidate ncRNAs on the basis that these are more likely to act in trans. We were therefore surprised to find that the GO term enrichment of adjacent genes for both CNSs and our candidate ncRNAs was similar. This cannot be explained by other factors, as random regions did not yield the same signal. One possibility is that the expression from these elements is of the type seen for some enhancer elements, where expression may open up genomic regions, making them accessible for cis-action. This phenomenon is well documented (Shen et al. 2018; Kim and Shiekhattar 2015), and our data suggest this may be prevalent across transcribed noncoding elements, thus blurring the boundaries between DNA-based cis-acting elements and trans-acting ncRNA genes.

Despite these apparent similarities, we nevertheless find distinct and distinctive epigenetic signatures that appear to delineate the expressed and unexpressed elements in our study. This suggests these may have distinct functions reflected in their distinct epigenetic signatures. These observations in turn suggest that it may be possible to use epigenetic signatures in the identification of functional expressed ncRNAs and CNSs. This may provide a major additional tool in noncoding element identification, particularly as it is difficult to confidently identify these without comparative genomic data. The comparative approach aids in building high-confidence annotations as sequence conservation is a strong signal of functional constraint. However, for species specific regulatory elements, which may be central to phenotypic change (King & Wilson), epigenetic signatures may help in the identification of newly-emerged noncoding elements.

In summary we identified 153475 conserved noncoding elements that are common in the 10 primate species used in the study. 59,870 elements were designated as ncRNA-gene-CEs and 93605 elements that showed no overlaps with annotated ncRNA genes as CNSs. CNSs showed higher signal strength for H3K4Me1 and H3K27ac which are histone modifications related with active enhancer regions compared to ncRNA-gene-CEs. Therefore we assume that many of these CNS elements might be functioning as active enhancers compared to primate common ncRNA-gene-CEs. It has been found that conserved elements are associated with active enhancer regions (Heintzman et al. 2009; Creyghton et al. 2010). Blum et al. 2012 found that 34% of their myoblast enhancers and 36% of myotube enhancers overlapped conserved noncoding sequences. It is currently known that even some of the ncRNA can also act as enhancer elements (Kim et al. 2010; Natoli and Andrau. 2012; Chen et al. 2017), but our primate common ncRNA-gene-CEs showed less association with active enhancer element modifications, signifying that these conserved elements may be related to other functions. Also we found that CNSs were enriched with histone modifications associated with regulation of development and active promoters. These findings with regards to CNSs are in congruence with already documented evidence (Inada et al. 2003; Bernat et al. 2006;Hettiarachchi et al. 2016).We found that primate common ncRNA-gene-CEs had significantly higher levels of H4K20Me1which is a histone modification associated with DNA repair than for developmental gene expression and regulation. This concept of RNA playing a role in DNA damage repair was not a widely explored concept. But currently there are several studies documenting that small and long ncRNA can play a role in DNA damage repair process (Sharma and Misteli. 2013; Michelini et al. 2017). Also D’Alessandro and Fagagna. 2018 reported that especially long ncRNA plays a role in genome stability. With regards to the primate common CNSs and ncRNA-gene-CEs we notice an interleaved functional arena with regards to gene regulation, whereas DNA damage repair is concerned we see that predominantly ncRNA play a key role, thus implying a slight differentiation in the functions related to genome stability. Also we found that primate common ncRNA-gene-CEs have a higher tendency to be located in open chromatin regions compared to rest of the genome, therefore these conserved ncRNA regions are easily accessible. This open conformation agrees with facilitating the regulatory functions that may be related to ncRNA.

We also found that primate common CNSs and ncRNA-gene-CEs are under purifying selection signifying that these regions have a functional constraint. Both types of CNSs showed lower frequency of derived alleles compared to random regions in human genome. This analysis was consistent across all three populations that we used in the study (Yoruba, Han Chinese and European). It has been known for a while that conserved non-coding sequences are under purifying selection and are not merely mutational cold spots (Drake et al. 2006; Katzman et al. 2007; Sakuraba et al. 2008). The primate common ncRNA-gene-CEs showed a higher percentage level of lower allele frequency compared to CNSs, which in a way signifies that the ncRNA-gene-CEs are under a strongly constrained selective pressure. It is expected that majority of conserved regions to be under purifying selection given the assumption that they have been conserved across long divergence times due to their functional importance, but our result here shows that even among conserved regions some of them can be under a stronger selective pressure which could be due indispensable functions that cannot be compromised in the genome.

Yet another interesting result of this study that is worth emphasizing is that we found primate common CNSs and ncRNA-gene-CEs have gone through a phase of accelerated evolution in several different stages during evolution before getting stabilized. It appears the faster mutation of sequences to adjust to new divergence events happen in waves. The human-gorilla common ancestor sequences showed 4 times faster evolutionary rate compared to human-gorilla-orangutan common ancestor for both ncRNA-gene-CEs and CNSs. This result suggests that certain mutational changes occurred at a faster rate in the human-gorilla common ancestor sequences to facilitate the functional requirements of the species after divergence. Also we observed that Hominoidae common ancestor showed accelerated evolution in CNS sequences after diverging from Hominoidae-old world monkey common ancestor. Also old world monkeys show about 2 fold acceleration after diverging from Hominoidae-old world monkey common ancestor. Going a step further from widely accepted belief that conserved regions are under constant purifying selection it is important to understand that some conserved regions change faster prior to getting stabilized and going through purifying selection to maintain what was achieved by accelerated mutation events. This scenario is possible where species after divergence adopt new set of features that may be physiological or morphological that cannot be regulated by already existing functional elements in a genome. Particularly Homininae also referred to as “African great apes” consisting human, chimpanzee and gorilla did particularly experience an increase in brain size and bipedalism (Lovejoy Co. 1980). Without enough evidence even though it is hard to point that the accelerated evolution in conserved regions might be responsible for special characteristics of this lineage, it has been found that certain CNSs are related to neuronal adhesion of human and chimpanzee (Prabakar et al. 2006).

It is known through experimental evidence that CNSs, actually do regulate the closest gene in most scenarios (Sumiyama et al. 2002; Bhatia et al. 2014). While considering the functional classification for the ncRNA-gene-CEs and CNSs we found that both groups are enriched with genes involved in regulating transcription and neuronal functions. Adopting the most widely accepted classical understanding this implies that ncRNA-gene-CEs and CNSs both actually might be responsible for similar functions. A recent study revealed that especially long ncRNA (lncRNA) being associated with development of central nervous system and neurodegenerative disorders (Wei et al. 2018). Despite the previous belief that lncRNA does not have a function and is a result of mistakes during transcription can now be challenged with computational and also some experimental evidence. This result further clarifies that CNSs that were identified as ncRNA-gene-CEs in this study might actually have the potential to be related to gene regulation prior any experimental evidence.

The GO for CNSs and ncRNA-gene-CEs showed evidence that likely target gene of these regions are enriched in brain and nervous system related functions. Even though this was a known fact for CNSs it wasn’t very clear how the ncRNA likely target genes behaved in genomes. Also further investigation showed that both CNSs and ncRNA-gene-CEs were related to brain and nervous system related tissues especially Amygdala, brain meninx, occipital lobe and spinal cord (fetal and adult both). Several studies document that CNSs are related to brain and nervous system tissues (Meyer et al. 2017). But our finding shows that ncRNA-gene-CEs related genes can also be related to nervous system development rather than house-keeping functions.

We found 40% of the primate common elements in mammalian common ancestor implying that about 60% of the primate common elements we identified may have solely originated in primate common ancestor and could be accounting for necessary functional requirements in the primate lineage. One interesting feature is that the newly originated elements also appear to predominantly cluster around genes related to central nervous system. This indirectly implies that further adoption of conserved elements was required for brain function or re-wiring of molecular pathways related to brain function in the primate lineage. One significant feature that should be noted is that there has been a gradual expansion in the neocortex volume in primates compared to insectivores and prosimians. Also further expansion in the primate lineage leading to, great apes having the largest neocortex (Stephen and Andy. 1970) in the animal kingdom. The neocortex is known as the centre for higher brain functions, such as decision making, reasoning, language and consciousness. The expansion of this region and functions related to it in primate lineage or more over higher level primates such as great apes may have required new regulatory mechanisms thus newly originated regulatory elements. Even though we cannot directly narrow down all the elements we have identified in this study are related to neocortex functions, the tissue expression analyses imply that the conserved elements here are related to brain regions compared to other organs.

Also hominoidae specific elements in this study being related to brain activity further solidify the reasoning above. Nevertheless it is important to note that mammalian common elements showed much higher functional fold enrichment for central nervous system related activity in comparison to both primate common and hominoidae specific elements. Could this mean that the determination of basic building blocks for proper functionality and structure of central nervous system occurred way before primates is something worth looking at in more detail?

In many instances showing similar pattern in target gene functionality being related to similar gene groups, potential enhancer region overrepresentation directly gives evidence that the primate common CNS and ncRNA-gene-CEs in our study have a tendency to function in a similar manner. Even though ncRNAs are presumably trans acting elements, what we have identified in our study shows that these ncRNA gene elements seem to resonate cis-acting CNSs.

Finally our findings in this study have shed light upon primate common, mammalian common and hominoidae specific ncRNA-gene-CEs and CNSs from numerous functional perspectives and have provided evidence that ncRNA-gene-CEs and CNSs both are important for proper functionality of genomes and ncRNA-gene conservation may also be indispensable to genomes than we give credit for.

## Declarations

### Availability of data material

The datasets produced during the current study are available from the corresponding author on reasonable request. All raw genomic data the author has used is publicly available and is being duly cited in the references.

### Contributions

Computational analyses / conceptual analyses based on results / Data interpretation / Methodology / writing of draft: Nilmini Hettiarachchi

## Ethics declarations

### Ethics approval and consent to participate

Not applicable

### Consent for publication

The author consents to the above manuscript being published in BMC Genomics.

## Acknowledgements

I specially thank Prof. A. Poole, Dr. M. Hoeppner who provided valuable constructive suggestions, views and proof reading support during this project. I also give my sincere thanks to others who provided suggestions.

