## Supplementary file 1 for "Primate deep conserved noncoding sequences and non-coding RNA: their possible relatedness to brain and Central Nervous System"

### Supplementary material

Supplementary Table 1– Number of histone modification signals located inside CNSs or ncRNA-gene-CEs

| Histone modification | Number of regions located inside CNSs | Number of regions located inside ncRNA-gene-CEs |
|---|---:|---:|
| H3K4Me1 | 144 | 100 |
| H3K4Me2 | 108 | 96 |
| H3K4Me3 | 79 | 53 |
| H3K9ac | 44 | 32 |
| H3K9Me3 | 44 | 20 |
| H3K27ac | 45 | 50 |
| H3K27Me3 | 36 | 23 |
| H3K36Me3 | 31 | 54 |
| H3K79Me2 | 9 | 9 |
| H4K20Me1 | 8 | 15 |

Supplementary Table 2– GO terms related  with closest genes for mammalian common CNSs and conserved ncRNA. (A)Gene Enrichment values for likely target genes for CNSs (B) Gene Enrichment values for likely target genes forncRNA-gene-CEs

(A)

| GO term | Fold enrichment | P-value |
|---|---|---|
| Regulation of chemotaxis | 3.19 | 1.62E-03 |
| Glutamate receptor signaling pathway | 2.77 | 4.60E-04 |
| Synaptic transmission | 2.70 | 4.61E-02 |
| Regulation of cell morphogenesis | 2.70 | 1.53E-03 |
| Negative regulation of signal transduction | 1.98 | 5.01E-04 |

(B)

| GO term | Fold enrichment | P-value |
|---|---|---|
| Regulation of synapse structure or activity | 4.32 | 9.50E-04 |
| Positive regulation of synaptic transmission | 3.28 | 1.17E-03 |
| Regulation of axogenesis | 3.14 | 1.05E-03 |
| Glutamate receptor signaling pathway | 2.95 | 4.05E-04 |
| Synaptic transmission | 2.89 | 2.09E-04 |

Supplementary Table 3– GO terms related with closest genes for hominoidae specific conserved elements. (A) overrepresented functional classifications of likely target genes (B) underrepresented GO terms

**(A)**

| GO term | Fold enrichment | P-value |
|---|---|---|
| Neuron development | 2.04 | 2.68E-05 |
| Neuron differentiation | 1.69 | 7.85E-04 |
| Chemical synaptic transmission | 1.58 | 5.12E-04 |
| Trans synaptic signaling | 1.58 | 5.12E-04 |
| Intracellular signal transduction | 1.42 | 6.03E-05 |

**(B)**

| GO term | Fold enrichment | P-value |
|---|---|---|
| Immune effector process | 0.31 | 2.05E-04 |
| Phagocytosis | 0.23 | 3.90E-04 |
| Defense response to bacterium | 0.06 | 1.30E-06 |
| Regulation of lymphocyte activation | <0.01 | 5.53E-06 |

Supplementary Table 4 **–** Broad feature categories of CNS-ncRNA cross homology sequences. This analysis took a total of 4772 cross homology elements and searched for coordinate overlaps in Ensembl feature categories.

| **Feature category** | **Numberof cross homology sequences overlapping a particular feature** |
|---|---|
| Histone related | 15 |
| TF binding related | 43 |
| miRNA target site | 449 |
| RNApolIII | 1 |

Supplementary Table 5 – CNSs and ncRNA-gene-CEs overlapping Ensembl verified enhancers, CTCF bindng sites and promoters.

| **Feature category** | **Number CNSs overalpping with regualtory feature** | **Number of ncRNA gene-CEs overlapping regualtory feature** |
|---|---|---|
| Enhancers | 5110 | 3403 |
| CTCF binding | 4189 | 2840 |
| Promoters | 4366 | 4632 |

Supplementary Table 6– GO terms related with closest genes for CTCF, enhancer and promoter overlapping mammalian common CNSs and conserved ncRNA. (A)Gene Enrichment values for likely target genes for CTCF-CNSs (B) Gene Enrichment values for likely target genes for CTCF ncRNA-gene-CEs (C)Gene Enrichment values for likely target genes for enhancer-CNSs (D) Gene Enrichment values for likely target genes for enhancer ncRNA-gene-CEs (E)Gene Enrichment values for likely target genes for promoter-CNSs (F) Gene Enrichment values for likely target genes for promoter ncRNA-gene-CEs

(A)

| GO term | Fold enrichment | P-value |
|---|---|---|
| Anterior posterior axon guidance | 7.66 | 9.49E-04 |
| positive regulation of axon extension involved in axon guidance | 7.66 | 9.49E-04 |
| netrin-activated signaling pathway | 7.15 | 1.80E-04 |
| positive regulation of axon guidance | 6.70 | 1.52E-03 |
| neuron projection extension involved in neuron projection guidance | 6.70 | 1.52E-03 |

(B)

| GO term | Fold enrichment | P-value |
|---|---|---|
| Cardiac muscle cell proliferation | 6.31 | 5.29E-04 |
| striated muscle cell proliferation | 5.89 | 7.27E-04 |
| muscle cell proliferation | 5.61 | 3.92E-04 |
| mammary gland lobule development | 4.81 | 8.84E-04 |
| mammary gland alveolus development | 4.81 | 8.84E-04 |

(C)

| GO term | Fold enrichment | P-value |
|---|---|---|
| trigeminal ganglion development | 10.74 | 7.81E-04 |
| nephric duct morphogenesis | 8.95 | 4.89E-06 |
| cranial ganglion development | 8.95 | 4.89E-06 |
| presynaptic active zone organization | 7.67 | 2.12E-03 |
| corticospinal tract morphogenesis | 7.67 | 2.12E-03 |

(D)

| GO term | Fold enrichment | P-value |
|---|---|---|
| pulmonary artery morphogenesis | 10.65 | 5.32E-04 |
| positive regulation of male gonad development | 8.28 | 1.21E-03 |
| positive regulation of extrinsic apoptotic signaling pathway in absence of ligand | 8.13 | 4.14E-04 |
| cardiac cell fate commitment | 8.13 | 4.14E-04 |
| nephric duct morphogenesis | 7.45 | 5.88E-04 |

(E)

| GO term | Fold enrichment | P-value |
|---|---|---|
| endothelial cell fate commitment | 8.74 | 2.13E-04 |
| bud elongation involved in lung branching | 7.49 | 1.06E-03 |
| epithelial cell fate commitment | 6.24 | 6.46E-05 |
| type B pancreatic cell development | 6.24 | 6.46E-05 |
| cell proliferation involved in kidney development | 6.12 | 9.15E-04 |

(F)

No statistically significant GO terms were found for ncRNA gene CEs overlapping promoter regions

Supplementary figure 1 - **Derived allele frequency for CNSs and ncRNA-gene-CEs with Han Chinese population.**(A) Derived Allele Frequencies for CNSs with Han Chinese population (B) Derived Allele Frequencies for ncRNA-gene-CEs with Han Chinese population data

(A)

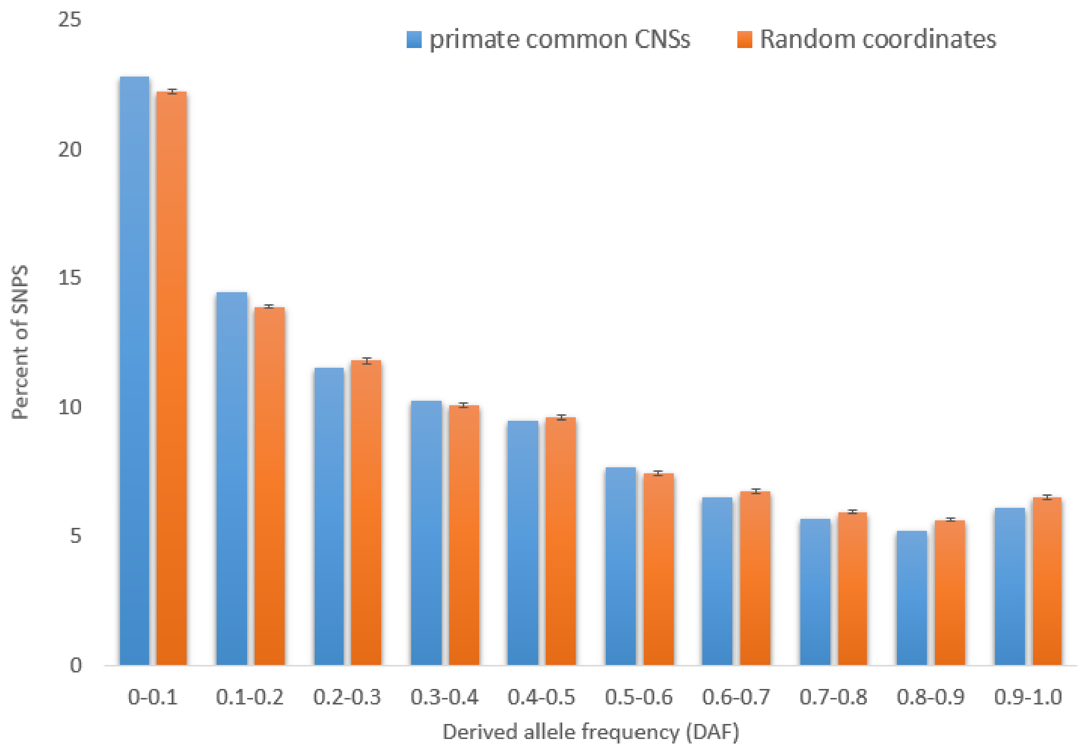


(B)

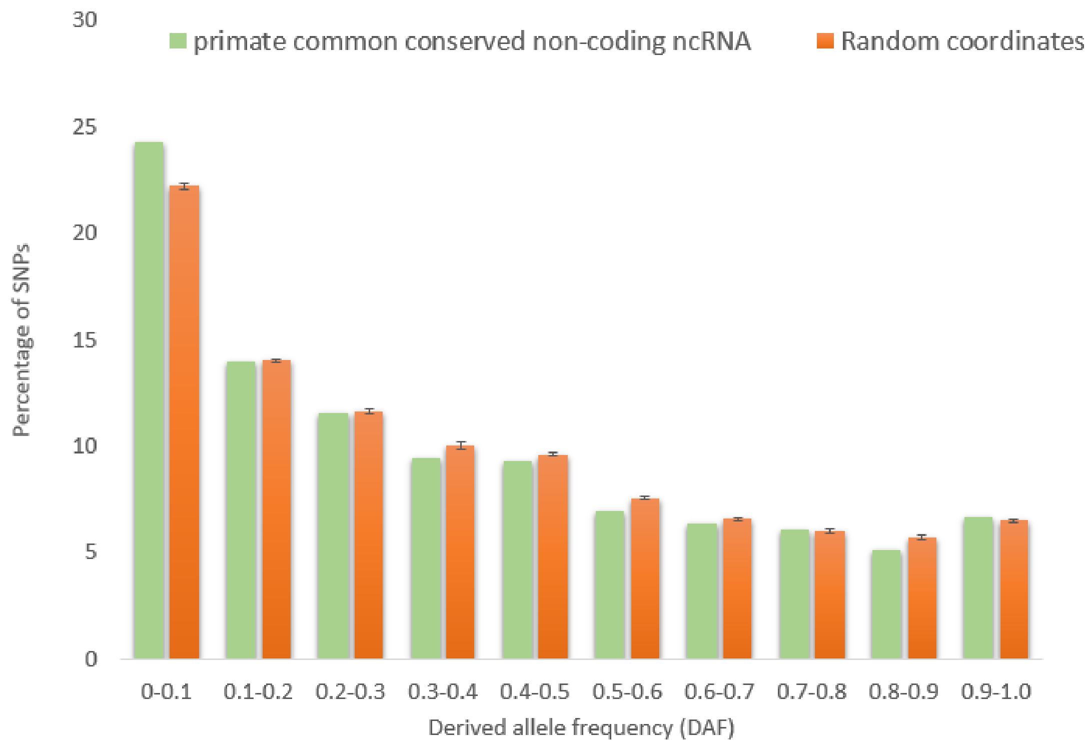

Supplementary Figure 2- **Derived allele frequency for CNSs and ncRNA-gene-CEs with European population.**(A) Derived Allele Frequencies for CNSs with European population (B) Derived Allele Frequencies for ncRNA-gene-CEs with European population data

(A)

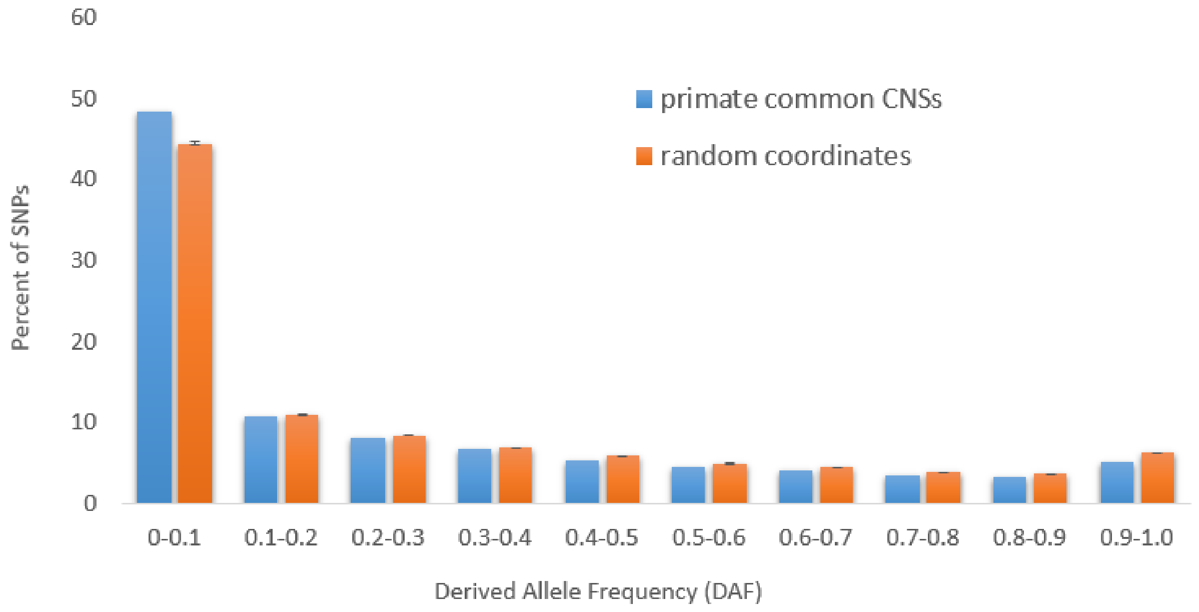


(B)

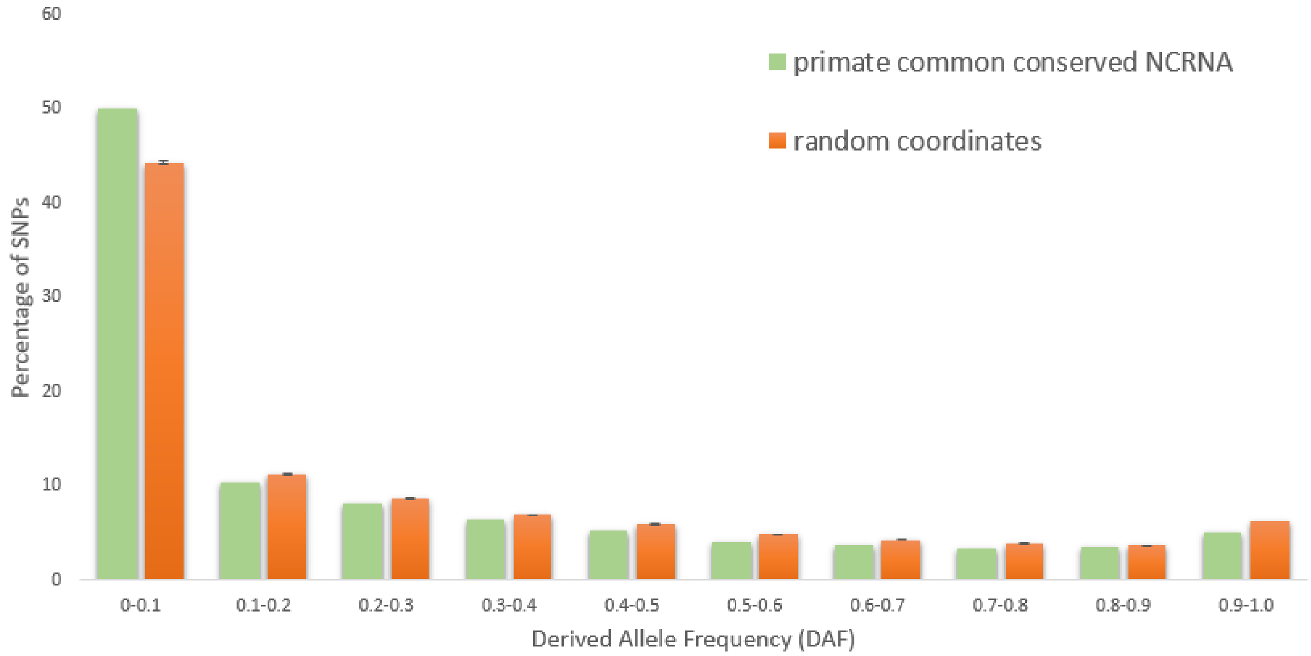

Supplementary Figure 3 – **Enhancer, CTCF binding and promoter elements in primate common CNSs and ncRNA-gene-CEs according to Ensembl regulatory track data.** (A) Enhancer elements overlapping CNSs, sampled CNSs, ncRNA-gene-CEs and random regions. (B) CTCF binding elements overlapping CNSs, sampled CNSs, ncRNA-gene-CEs and random regions. (C) Promoter elements overlapping CNSs, sampled CNSs, ncRNA-gene-CEs and random regions. Statistically significant (p <0.05) combinations are depicted by * and non-significance by NS.

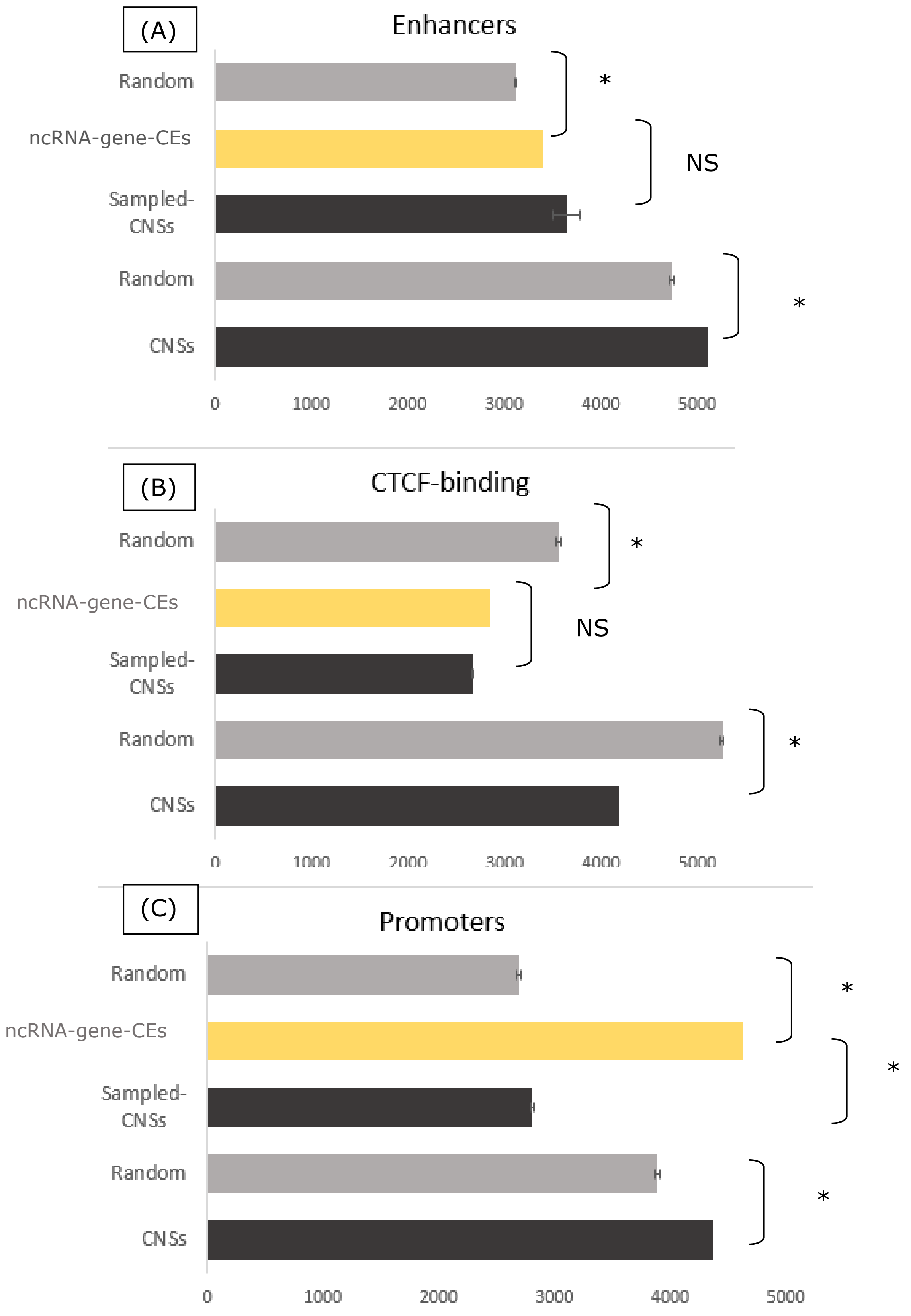

Supplementary Figure 4- **percentage occurrence of gene-CNS gene-ncRNA clusters of primate common conserved noncoding regions.**The figure colour legend represents all cases of gene-CNS or ncRNA associations. i.e1gene - 1CNS/1 ncRNA, 1 gene-multiple CNSs/ncRNA associations and percentage occurrence.

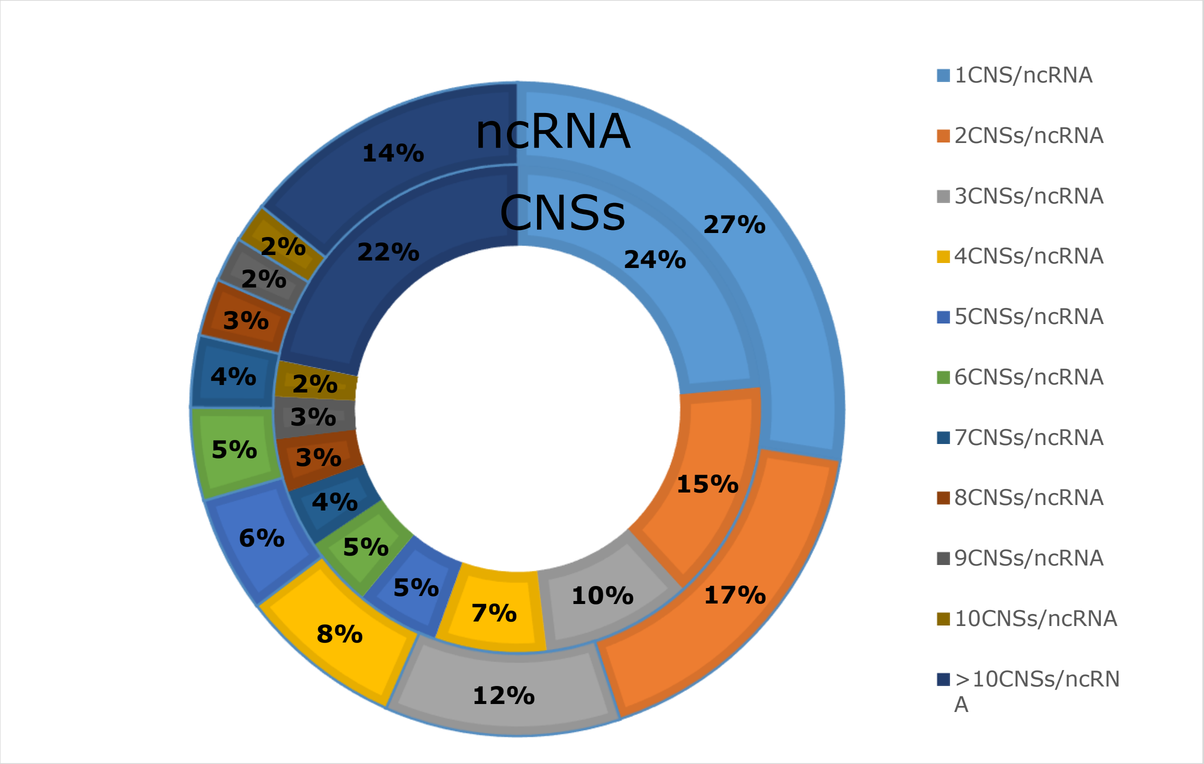
